# Genetics and resource availability shape divergence in life history and behavior between locally-adapted populations of Atlantic mollies (*Poecilia mexicana*, Poeciliidae)

**DOI:** 10.1101/2022.11.23.517737

**Authors:** John L. Coffin, Bethany L. Williams, Michael Tobler

## Abstract

Phenotypic variation is common along environmental gradients, but it is often unknown to what extent it results from genetic differentiation between populations or phenotypic plasticity. We studied populations of a livebearing fish that have colonized streams rich in toxic hydrogen sulfide (H_2_S). In nature, there is strong phenotypic differentiation between adjacent sulfidic and nonsulfidic populations. In this study, we varied food availability to pregnant mothers from different populations to induce maternal effects, a form of plasticity, and repeatedly measured life-history and behavioral traits throughout the offspring’s ontogeny. Genetic differentiation affected most of the traits we measured, as sulfidic offspring tended to be born larger, mature later, have lower burst swimming performance, be more exploratory, and feed less accurately. In contrast, maternal effects impacted few traits and at a smaller magnitude, even though offspring from poorly provisioned mothers tended to be born larger and be more exploratory. Population differences and maternal effects (when both were present) acted synergistically, and there was no evidence for population differences in plasticity. Overall, our study suggests that phenotypic divergence between these populations in nature is primarily caused by genetic differentiation, and that plasticity mediated by maternal effects accentuates—but does not cause—differences between populations.

## Introduction

Phenotypic variation is at the heart of evolutionary analyses because it links the cause (natural selection) to the consequence (genotypic change) of adaptive evolution (Lande & Arnold, 1983). We have known how inheritance causes resemblance between parents and their offspring for well over a century, reflecting a genetic component to phenotypic variation (Stenseth *et al*. 2022). However, trait variation can also be influenced by phenotypic plasticity, where a single genotype can give rise to alternate phenotypes in response to internal or environmental cues (West-Eberhard, 1989; Pigliucci, 2001). In addition, the environment experienced by parents can affect phenotypes of their offspring (*i.e*., parental effects; Uller 2008; Badyaev and Uller 2009), representing a case of plasticity that spans generational boundaries. Phenotypic variation in nature can therefore arise from genetic differences among individuals, plasticity induced by individual exposure to different environmental conditions, plasticity induced by parental effects, and their interactions (Scheiner 1993; Dingemanse and Araya-Ajoy 2015). For many natural systems, we know little about the origins of phenotypic variation, even though it critically shapes our inference of adaptation in natural populations.

Plasticity induced by parental effects is particularly strong from mothers due to their higher reproductive investment and, in viviparous species, the physically intimate relationship with their developing young (Lindholm, Hunt, & Brooks, 2006; Wolf & Wade, 2009). Such maternal effects are widespread in nature (Mousseau and Fox 1998) and can impact trait expression and evolution (Rossiter, 1996; Wilson *et al*., 2005; Beckerman *et al*., 2006). Maternal effects can be adaptive if the expression of offspring traits is biased to match the environment experienced by the mother (Marshall and Uller 2007), or if mothers in good condition are able to endow phenotypes that provide a competitive advantage to their offspring in any environment (Grafen, 1988; Monaghan, 2008; Van Allen *et al*., 2021). However, maternal effects can also be maladaptive and produce mismatches between offspring phenotype and environment, as documented in some organisms responding to anthropogenic climate change (Schuler and Orrock 2012; Leonard and Lancaster 2020). Maladaptive maternal effects can also be related to stress, where physiological stress responses in mothers have unintended negative side effects on offspring (MacLeod, While, & Uller, 2021). Regardless of whether maternal effects are adaptive, they are important biological phenomena that warrant careful attention and explicit accounting in evolutionary analyses due to the non-genetic effects on phenotypic expression.

Phenotypic variation in nature is common along environmental gradients, but it is often unclear whether it is caused by genetic differentiation among populations or plastic effects that arise from population-specific environmental exposure histories experienced by mothers or directly by their offspring. For example, freshwater springs rich in toxic hydrogen sulfide (H_2_S) in the Grijalva River basin of southern Mexico are extreme environments that are connected to adjacent nontoxic streams, and stark phenotypic gradients can be observed in fish occupying these habitats in as little as a few meters. Sulfide springs are complex ecosystems with several correlated sources of selection (Tobler *et al*., 2016b). H_2_S is toxic because it disrupts aerobic ATP production (Reiffenstein, Hulbert, & Roth, 1992; Cooper & Brown, 2008), but habitats rich in H_2_S also differ from nonsulfidic habitats in other physical and chemical water parameters (dissolved oxygen concentrations, pH, and salinity), and the presence of competitors and predators (Riesch *et al*., 2010a; Tobler *et al*., 2011; Greenway et al., 2014). Resource availability also differs greatly between habitat types; fish in nonsulfidic environments eat primarily algae and detritus, while fish in sulfidic environments have shifted to eating primarily sulfide bacteria and invertebrates (Tobler *et al*., 2015). Populations of Atlantic mollies (Poecilia mexicana), a species of livebearing fish of the family Poeciliidae, have independently colonized and adapted to sulfidic streams across multiple river drainages, and previous studies have documented that colonization of sulfide springs has been associated with convergent changes in morphology, locomotion, and respiration (Tobler & Hastings, 2011; Camarillo, Arias Rodriguez, & Tobler, 2020), behavior (Plath *et al*., 2007; Lukas *et al*., 2021; Doran *et al*., 2022), physiology (Tobler et al., 2011; Barts *et al*., 2018; Greenway *et al*., 2020), and life history traits (Riesch *et al*., 2011a; Riesch, Plath, & Schlupp, 2011b; Riesch *et al*., 2014). Phenotypic divergence between sulfidic and nonsulfidic mollies likely has a significant genetic component, because it coincides with strong genetic differentiation between populations, even though there are no physical barriers separating populations in the different habitat types (Palacios *et al*., 2013; Plath *et al*., 2013; Riesch *et al*., 2016b). However, trait variation between populations likely also has an environmental component; while population differentiation persists in captive populations reared in common garden conditions in the lab (Tobler *et al*., 2016a; Greenway *et al*., 2020), there is also evidence for plasticity caused by short-term exposure to different environmental conditions (Bierbach *et al*., 2011; Passow *et al*., 2017a). In addition, the impact of maternal effects on offspring trait expression remains to be investigated in these livebearing fish. Hence, we tested how genetic differentiation, maternal effects, and their interactions shape phenotypic expression in *P. mexicana* populations from sulfidic and nonsulfidic habitats. To induce maternal effects, we manipulated resource availability to pregnant mothers, because natural populations vary substantially in nutritional state. Fish in sulfidic habitats are consistently under food stress, exhibiting significantly reduced body condition (both when inferred through length-weight regression and body fat content analysis; Plath *et al*., 2005; Tobler *et al*., 2006; Tobler, 2008). Food stress arises as a consequence of constraints associated with resource acquisition; because H_2_S coincides with and exacerbates hypoxia, fish from sulfidic habitats have to trade off performing aquatic surface respiration, a compensatory behavior to access better-oxygenated surface waters, with benthic foraging (Tobler *et al*., 2009). Accordingly, populations in sulfide springs have adaptations to low resource availability, including reductions in routine metabolic rates and energetically expensive tissues like the brain (Schulz-Mirbach *et al*., 2016; Passow, Arias-Rodriguez, & Tobler, 2017b). In other species, including some poeciliids, resource availability experienced by mothers has been shown to impact offspring trait expression (Reznick, Callahan, & Llauredo, 1996a; Altmann & Alberts, 2005; Boots & Roberts, 2012), and different population histories in terms of exposure to food stress in *P. mexicana* may have caused changes in resource-induced maternal effects.

To quantify population differentiation and maternal effects in populations of *P. mexicana*, we followed families of offspring from birth to the onset of maturation and repeatedly quantified a host of complex phenotypic traits. Focal traits included brood size, size at birth, and age at maturity as well as ontogenetic trajectories in growth rates, burst swimming, exploratory behavior, and feeding accuracy. We chose these traits because they likely affect fitness, and we have prior knowledge for many of them from natural populations, providing us with a framework to make a *priori* predictions. Specifically, our experiments sought to address four specific questions: (1) Is there evidence for phenotypic trait differences between populations from sulfidic and nonsulfidic habitats that persist in fish reared in a common-garden environment for multiple generations? Divergence in phenotypic traits between populations regardless of maternal food treatments would indicate that trait differentiation is due to genetic variation between populations. (2) Is there evidence for maternal effects in response to resource availability? Significant differences in offspring traits between maternal food treatments, irrespective of population of origin, would suggest resource-induced maternal effects. (3) How do functional traits vary throughout ontogeny, and how do population differences and maternal effects interact with ontogeny? Age is a major determinant in the expression of many traits (Hegyi *et al*., 2006; Zhang *et al*., 2015), but how population differences and maternal effects impact trait expression through ontogeny is less clear. In other poeciliids, maternal effects tend to be present at birth and decline with age (Lindholm *et al*., 2006). In contrast, population differences between sulfidic and nonsulfidic *P. mexicana* are stark in adults, suggesting differences may emerge early in life and even increase throughout ontogeny (Riesch *et al*., 2011a). Accordingly, we predicted that age would impact most of the traits measured, that population differentiation would increase with age, and that maternal effects would diminish with age. (4) How do population differences interact with maternal effects? A difference in how each population responds to variation in maternal resource availability would indicate genotype-by-environment interactions. Fish from sulfidic habitats generally face constraints in resource levels, while those from nonsulfidic habitats have access to more abundant resources (Tobler *et al*., 2006, 2009; Tobler, 2008). Hence, we predicted that trait variation induced by the low-food treatment would occur in the same direction as trait variation produced by differences between the nonsulfidic and sulfidic populations (*i.e*., maternal effects would be aligned with population differences). In this case, maternal effects would accentuate divergence between populations that resembles patterns of variation found in the wild. Alternatively, low-resource traits may be canalized in the sulfidic population because sulfidic individuals are constantly food stressed in nature. In this case, maternal effects may be weaker in the sulfidic population.

## Methods

### Experimental overview

For our experiments, we used two lab-reared populations of *Poecilia mexicana* originating from wild-caught relatives in the Tacotalpa River drainage. One population originated from a sulfide spring complex called El Azufre I (according to Plath *et al*., 2013; hereafter referred to as the sulfidic population or ecotype), and the other population was from a nonsulfidic stream connected to the mainstem of the Tacotalpa River, called Arroyo Bonita (according to Plath *et al*., 2013; hereafter referred to as the nonsulfidic population or ecotype). Both populations were reared in 680-liter stock tanks filled with filtered tap water. Tanks were fed *ad libitum* twice daily with commercial dry fish food (Purina), and ∼50% of the water was exchanged weekly. Mothers used in this experiment were raised under common-garden conditions for at least three generations (*i.e*., they were at least great-grandchildren from individuals originally collected in the wild).

From each tank, 30 females were caught with a dipnet and isolated in a 20-liter tank with an aerating filter and a bundle of plastic mesh as shelter for newborn fry. Females were fed twice daily, once with aquatic gel diet for omnivorous fish (Mazuri) and once with freshly hatched *Artemia* nauplii (Brine Shrimp Direct). Females were randomly assigned to either a “high-food” diet (0.32 ml per feeding) or a “low-food” diet (0.08 ml per feeding), with equal numbers of females from each population being assigned to each group (Figure 1). Each tank was checked daily for newborn fry. By the end of the experiment, 18 sulfidic females gave birth (12 from the high food treatment and 6 from the low food treatment) and 22 nonsulfidic females produced a brood (12 from the high food treatment and 10 from the low food treatment). The amount of time each female was in the food treatment before giving birth varied (mean 28.2 days, range 0-99 days). As some females gave birth during the acclimation period or immediately after beginning the food treatments, we included treatment length as a potential covariate in all analytical models. Additionally, for a subset of models we tested whether using females that were in the treatment for at least seven days would affect the resulting top models (Table S1). Because the models were largely unaffected, we chose to include all broods to increase statistical power to detect population differences throughout ontogeny.

**Figure 1.**
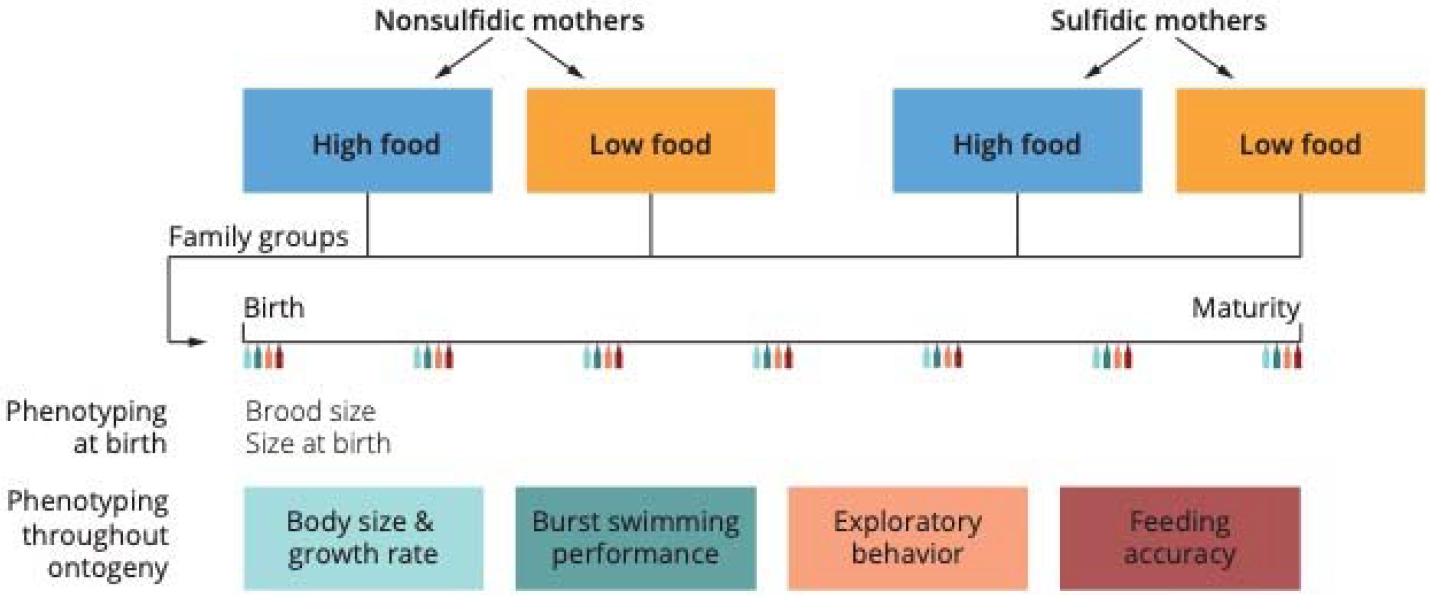
Overview of our experimental design. We subjected pregnant sulfidic and nonsulfidic mothers to either a high- or a low-food treatment and measured seven traits in their offspring throughout their development. These traits include four life history traits (birth size, brood size, growth rate, and age at maturity) and three behavioral traits (burst swimming, exploratory behavior, and feeding accuracy).

Whenever a brood was born, we recorded the brood size, date of birth for each family, the number of days that the female was in the food treatment, and the mother’s standard length (distance from the anterior tip of the snout to the posterior end of the caudal peduncle, mm). Mothers were then removed from the tanks, and we randomly selected 15 newborn fry (if available) from each family to remain in the experimental tanks to minimize density-dependent effects.

From that point on, we followed the developing families through ontogeny. All fry—irrespective of the mother’s food treatment—received the same amount of food; they were fed *ad libitum* twice daily with a mixture of decapsulated brine shrimp eggs (Brine Shrimp Direct) and dry food. We assessed offspring phenotypes at approximately weekly intervals by measuring life history traits (size at birth, weekly growth rate, and age at maturity) and behavioral traits (burst swimming, exploratory behavior, and feeding accuracy; see Figure 1). We specifically tested whether there were differences in these phenotypes between maternal food treatments (*i.e*., maternal effects), between populations (*i.e*., sulfidic vs. nonsulfidic, indicating effects of evolved population differences), or both. All analyses were conducted in R 4.0.5 (R Core Team, 2023). Code and data to reproduce all analyses can be found on GitHub (https://github.com/michitobler/common-garden).

### Size at birth and growth rate

To measure size at birth, we photographed each family from above on the day of their birth with a Nikon D90 digital camera fitted with an AF-S Micro NIKKOR 105mm f/2.8 lens. A ruler was included in the background of each image. Images were imported to ImageJ v. 1.53 (Schneider, Rasband, & Eliceiri, 2012) and calibrated by setting the scale. We measured the standard length (mm) of each offspring in the family. These measurements were averaged across all individuals to obtain a single mean size at birth for each family. This measurement was completed weekly, and the measurement for each family was subtracted from that family’s measurement from the week prior to obtain an average growth rate [mm/week]. To account for allometric differences in growth rate, we converted the average weekly growth rates to a proportional growth rate by dividing the average weekly growth rate by the mean body size the week prior.

### Age at maturity

We estimated the minimum age at maturity for each family using morphological characteristics of male sexual maturity. While the sexes of juvenile livebearers are difficult to distinguish, at the onset of sexual maturity, the male anal fin is modified into an intromittent organ (gonopodium), while it remains unmodified in females (Rosen & Gordon, 1953; Chambers, 1987). We therefore measured minimum age at maturity (days) as the time it took to for the first male in a group to develop its complete gonopodium as judged by the presence of a fleshy palp on anal fin ray 3.

### Burst swimming

Most fish avoid predation with a highly conserved, reflexive escape response that causes the head to move away from the stimulus, bending the body into a ‘C’ shape (Eaton, Bombardieri, & Meyer, 1977). Then, a strong stroke of the caudal fin starts the movement away from the stimulus (Domenici & Blake, 1997). This process (bending into a ‘C’ and propelling away from the stimulus) is known as a C-start response and is frequently used as a metric of escape performance in fish (Walker, 1997; Ghalambor, Reznick, & Walker, 2004; Langerhans *et al*., 2004; Camarillo *et al*., 2020). To quantify this burst swimming behavior, we adopted the methods and metrics used by prior studies (Langerhans *et al*., 2004; Ingley *et al*., 2016; Camarillo *et al*., 2020). We placed a haphazardly chosen individual from each family in a glass petri dish (9 cm in diameter, containing 2 cm of water) with opaque sides suspended above an angled mirror, providing a ventral view of each fish. After 5 min of acclimation, we struck the surface of the water within a body length of the fish with a probe and recorded the movement of the fish from below with a Sony NEX-FS700R camcorder at 60 frames per second (fps) and 1080p resolution. We converted the resulting .mts files into .mp4 files (to enhance compatibility with downstream applications) with FFmpeg (Tomar, 2006).

We used DLTdv8 (Hedrick, 2008) to digitize the 2-D location of the isthmus (*i.e*., the area on the ventral surface of the head where the opercula converge) of the fish in each frame. Digitized points were then used to calculate the maximum velocity (*v*_max_, mm/s), maximum acceleration (*a*_max_, mm/s^2^), and net distance traveled (*d*_net_, mm of displacement within 1/12^th^ of a second after the C-start). To calculate *v*_max_, we calculated the straight-line distance between each pair of successive digitized points, divided this distance by the inverse of the frame rate (60 fps), and found the maximum value between any two points. *a*_max_ was calculated by subtracting the velocity value at each point from the velocity value at the point immediately preceding it and finding the maximum value. *d*_net_ was calculated by recording the 2-D position immediately after the fish ends the C-start with a single stroke of the caudal fin, and then recording the 2-D position 1/12^th^ of a second later (5 frames later) and calculating the straight-line distance between the two points. To reduce the dimensionality of this dataset, we conducted a principal component analysis (PCA) using the prcomp() function with a correlation matrix. There was one principal component with an eigenvalue greater than 1 (explaining 85.6 % of the total variance), which was retained as a compound metric of burst swimming performance. Positive scores along this principal component axis were associated with higher velocity, acceleration, and distance traveled (Table S2A). All mathematical operations were conducted using packages contained in the base distribution of R.

### Exploratory behavior

We used an open field test to quantify the exploratory tendencies of fry. We filled a Styrofoam cup (9 cm in diameter) with 3 cm of water and covered the arena with a sheet of glass. We haphazardly selected one individual from each family and placed it in the arena undisturbed for a 5-min acclimation period, after which we recorded 5 min of video from above with a GoPro Hero 4 (1080p resolution, linear field of view, 30 fps).

We used idTracker (v. 2.1, bundled with 64-bit Matlab Compiler Runtime 8.3; Pérez-Escudero *et al*., 2014) to track the 2-D location of the fish automatically for the entire 5-min recording. We set the number of individuals to 1 and manually determined the intensity threshold (0.5–0.8) and minimum size (40–250 pixels) for each video. We also imported a still frame from each video into ImageJ to measure the centroid coordinates and arena radius.

Using the 2-D coordinates in each frame, we calculated several metrics of motion that we used as proxies for exploratory behavior. We calculated distance traveled between each pair of successive points, the velocity, the acceleration, and the total cumulative distance traveled (*d*_total_, mm), as described above. We also calculated average velocity (*v*_avg_, mm/s), maximum velocity (*v*_max_, mm/s), and maximum acceleration (*a*_max_, mm/s^2^) by finding the means and maxima of all velocity and acceleration values. Finally, we calculated the proportional average distance from the center of the arena (*d*_center_, dimensionless) by calculating the distance from the fish’s location to the centroid of the arena and dividing this value by the arena radius. Videos were excluded if the fish was completely still in all frames. To reduce the dimensionality of and observe the correlation structure within this dataset, we ran a PCA on *v*_avg_, *v*_max_, *a*_max_, *d*_total_, and *d*_center_ with a correlation matrix. We retained scores along the first PC axis (explaining 56.9 % of the total variance) as a composite exploratory behavior score. As shown in Table S2B, higher PC1 scores were associated with more exploratory behavior (positive loadings for all variables).

### Feeding accuracy

To measure feeding accuracy, we withheld food from individuals of each tank for 24 h prior to the experiment and placed one haphazardly selected fry in a viewing tank for a 5-min acclimation period. The viewing tank was constructed by connecting two 10 cm × 10 cm sheets of glass to either side of 1 cm diameter glass rods with silicone. The rear wall of the viewing tank was covered with a black sheet of plastic to enhance contrast between the background and the fish in the tank. The feeding solution consisted of 1 g of freshly hatched, live *Artemia* nauplii diluted into 100 ml of filtered tap water. After 5-min of acclimation, we added 0.08 ml of the feeding solution to the viewing tank and recorded 5 min of video with the camcorder. We analyzed the video frame-by-frame using BORIS v. 7.13 (Friard & Gamba, 2016) and recorded the number of successful strikes (a feeding strike that ends in consumption of the food item) and unsuccessful strikes (a feeding strike that either misses the food item or results in regurgitation of the food item). We divided the number of successful strikes by the total number of strikes to get an overall proportional estimate of feeding accuracy, which was arcsine-square-root transformed prior to analysis.

### Statistical analyses of individual traits

There were many potential sources of variation in our experiment. Other than the effects of interest for our study (population, maternal food treatment, and their interaction), the observed variation in traits could also have arisen from differences in fry age, maternal body size (standard length), the duration of her food treatment, and brood size. Consequently, we used a model selection approach to find the models that were best supported by our data for each experiment separately. For each phenotype, we created a global model that contained all possible effects. For phenotypes that were only measured once for each family (size at birth, brood size, and age at maturity), we used a general linear model using the lm() function from the STATS package v. 3.6.2, and for phenotypes that were measured repeatedly through development (growth rate, burst swimming, exploratory behavior, and feeding accuracy), we used a linear mixed model implemented with the lmer() function from the LME 4 package v. 1.1-26 (Bates *et al*., 2015) that included ‘family’ as a random effect. We then used the dredge() function from the M_U_MI_N_ package v. 1.47.1 (Bartón 2009) to create a model selection table based on the effects contained in the global model, with different models ranked and weighted based on AIC_c_ (Johnson & Omland, 2004). Full model selection tables are available for each phenotype in Table S3. To avoid overfitting, we limited the models to a maximum of four terms. We chose the top-supported model for each trait and analyzed it with either type-III analysis of variance (ANOVA) or Wald’s chi-square tests (see Table 1). Quantification and visualization of effects was accomplished by calculating and plotting estimated marginal means for the effects of ‘Population’ and/or ‘Food Treatment’, depending on the best-supported model, using the Effect() function from the EFFECTS package v. 4.2-2 (Fox & Weisberg, 2018a,b). To aid in drawing inferences from our models, we generated 95% confidence intervals (CI) for model coefficients in our top models using the confint() function form the base R distribution. Effect sizes were calculated as partial eta-squared (***η***_p_^2^), which represents the proportion of variance explained by a particular variable after accounting for the variance explained by all other variables. We calculated ***η***_p_^2^ with the etasq() function (Fox *et al*. 2021) for general linear models or the eta_squared() function (Ben-Shachar, Lüdecke, & Makowski, 2020) for linear mixed models.

**Table 1.** Model terms in the best-supported model for each dependent variable. Empty cells represent terms that were not present in the best-supported model for that dependent variable. Type-III ANOVA or Wald’s Chi-square tests were used to determine whether model terms were significantly different from zero. F-statistics or *χ*^2^ statistics are presented for each included model term, with the degrees of freedom in a subscript. P-values are presented underneath each test statistic.

| Dependent Variable | Brood Size | Treatment Length | Mother SL | Age | Population | Food Treatment |
| --- | --- | --- | --- | --- | --- | --- |
| Size at Birth | $F_{1,35} = 20.876$<br>$P < 0.001$ | | $F_{1,35} = 16.593$<br>$P < 0.001$ | | $F_{1,35} = 20.032$<br>$P < 0.001$ | $F_{1,35} = 4.354$<br>$P = 0.044$ |
| Brood Size | | $F_{1,37} = 8.926$<br>$P = 0.005$ | $F_{1,37} = 23.342$<br>$P < 0.001$ | | | |
| Age at Maturity | | | | | $F_{1,26} = 4.420$<br>$P = 0.045$ | |
| Growth Rate at Birth | | | | | $F_{1,37} = 5.025$<br>$P = 0.031$ | |
| Overall Growth Rate | | | | $\chi^2_1 = 147.91$<br>$P < 0.001$ | | |
| Burst Swimming at Birth | | | $F_{1,39} = 5.025$<br>$P = 0.121$ | | | |
| Overall Burst Swimming | | | | $\chi^2_1 = 30.178$<br>$P < 0.001$ | $\chi^2_1 = 4.308$<br>$P = 0.038$ | |
| Exploratory Behavior at Birth | | | | | | $F_{1,31} = 2.727$<br>$P = 0.108$ |
| Overall Exploratory Behavior | | | | | $\chi^2_1 = 7.513$<br>$P = 0.006$ | $\chi^2_1 = 4.245$<br>$P = 0.039$ |
| Feeding Accuracy at Birth | | $F_{1,38} = 0.775$<br>$P = 0.384$ | | | | |
| Overall Feeding Accuracy | | | | | $\chi^2_1 = 4.645$<br>$P = 0.031$ | |
| Multivariate Trait Variation PC1 | | $F_{1,24} = 15.817$<br>$P < 0.001$ | | | $F_{1,24} = 14.984$<br>$P < 0.001$ | $F_{1,24} = 4.746$<br>$P = 0.039$ |
| Multivariate Trait Variation PC2 | | $F_{1,24} = 7.722$<br>$P = 0.010$ | $F_{1,24} = 5.292$<br>$P = 0.030$ | | $F_{1,24} = 3.321$<br>$P = 0.081$ | |

### Multivariate analysis

Since selection ultimately acts on complex, multivariate phenotypes (Lande & Arnold, 1983), we sought to understand how the traits measured for our analyses vary and covary to jointly shape multivariate phenotypes. To do so, we averaged each phenotype across all ages for each tank. We used this approach rather than including age as a covariate and analyzing raw phenotypic scores because of timing and logistical constraints that made it impossible to conduct each experiment on offspring that were exactly the same age. We analyzed the averaged phenotypes with a PCA (correlation matrix) and used scores along the first two principal components—which had eigenvalues greater than 1—as dependent variables. We analyzed principal component scores along each axis separately because the axes, by definition, were orthogonal. For each axis, we created a global model containing all possible effects and selected the best-supported model based on AIC_c_, as explained above.

## Results

We measured seven functional traits in offspring from a sulfidic and a nonsulfidic population of *P. mexicana* throughout ontogeny. Model selection tables for the analysis of each trait at birth and separately across all ages can be found in Table S3, and the best-supported models are summarized in Table 1. For brevity, we will present results in the context of our hypotheses outlined in the Introduction, focusing on how all traits vary through ontogeny in terms of population differentiation, maternal effects, and their interaction, rather than presenting the results for each trait separately.

### Is there evidence for population differentiation?

We found evidence for significant population differences in five of the seven traits measured: size at birth, age at maturity, burst swimming, exploratory behavior, and feeding accuracy (but not brood size or growth rate). Age at maturity, burst swimming, and feeding accuracy were significantly different between populations, but not between food treatments, whereas there were significant effects of ‘Population’ and ‘Food Treatment’ (but no interactions) on size at birth and exploratory behavior (see Table 1 and below). Sulfidic individuals were born 9.2 % larger [*F*_1,35_ = 20.032; *P* < 0.001; estimated marginal mean for sulfidic fish (EMM_S_) = 10.03 mm, estimated marginal mean for nonsulfidic fish (EMM_NS_) = 9.18 mm; ***η***_p_^2^ = 0.36; Figure 2A], matured an average of 10.5 days later (*F*_1.26_= 4.420; *P* = 0.045; EMM_S_ = 54.2 days, EMM_NS_ = 43.7 days; ***η***_p_^2^ = 0.15), had significantly lower burst swimming performance (*χ*^2^ = 4.308; *P* = 0.038; CI: –0.94, –0.03; ***η***_p_ ^2^ = 0.03), were more exploratory (*χ*^2^_1_ = 7.513; *P* = 0.006; CI: 0.29, 1.70; ***η***_p_^2^ = 0.28), and were 7.2% less successful during feeding (*χ*^2^_1_ = 4.645; *P* = 0.031; EMM_S_ = 0.72, EMM_NS_ = 0.80; ***η***_p_^2^ = 0.04; Figure 3).

**Figure 2.**
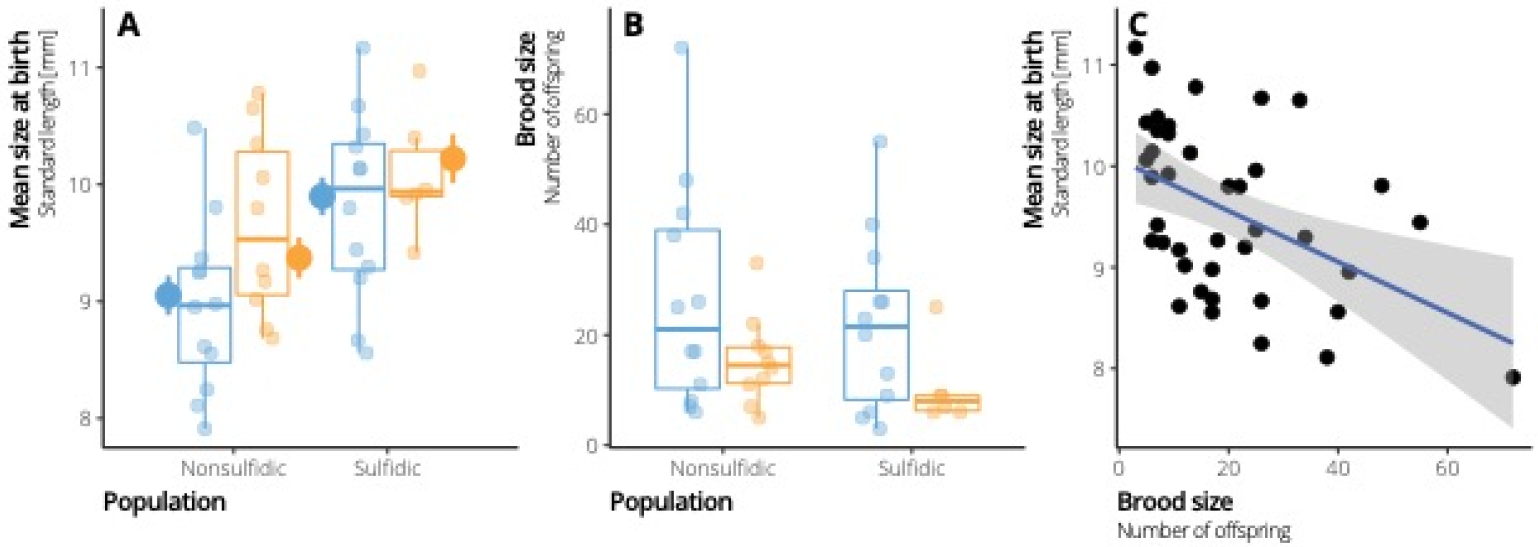
Plots of average standard length (A), brood size (B), and their relationship (C) for each family. Data is summarized in boxplots with raw data points overlaid. In both populations, the high-food treatment is shown in blue, and the low-food treatment is shown in orange. When the best-supported model for a phenotype contained the terms ‘Population’ and ‘Food Treatment’, the estimated marginal means for those effects were visualized as large points.

**Figure 3.**
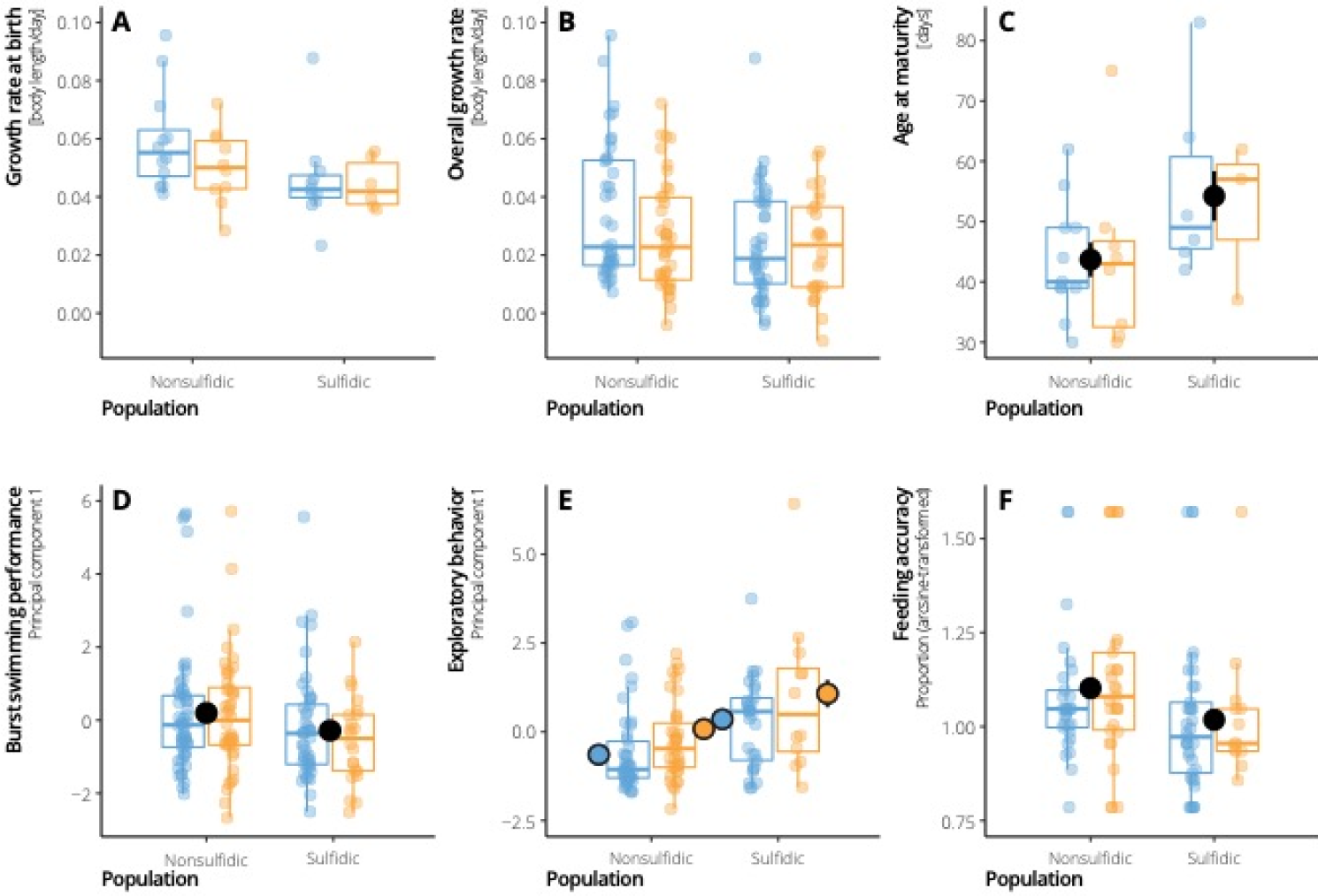
Data for each individual phenotype, including (A) growth rate at birth, (B) overall growth rate, (C) age at maturity, (D) burst swimming, (E) exploratory behavior, and (F) feeding accuracy. Data is summarized in boxplots with raw data points overlaid. In both populations, the high-food treatment is shown in blue, and the low-food treatment is shown in orange. When the best-supported model for a phenotype contained the terms ‘Population’ and/or ‘Food Treatment’, the estimated marginal means for those effects were visualized as large points.

### Is there evidence for maternal effects in response to resource availability?

To test whether maternal effects induced by resource availability during pregnancy impact functional traits in offspring, we compared each phenotype between offspring born to mothers who experienced a high-food environment and mothers who experienced a low-food environment. Mothers in low-food treatments produced offspring that were 3.4% larger at birth [*F*_1,35_ = 4.354; *P* = 0.044; estimated marginal mean for fish in the low-food treatment (EMM_low_) = 9.8 mm, estimated marginal mean for fish in the high-food treatment (EMM_high_) = 9.4 mm; ***η***_p_^2^ = 0.11; Figure 2A] and were more exploratory (*χ*^2^_1_ = 4.245; *P* = 0.039; CI: –0.04, –1.42; ***η***_p_^2^ = 0.18; Figure 3E). For all remaining traits (brood size, age at maturity, growth rate, burst swimming, and feeding accuracy), ‘Food Treatment’ was not included in the best-supported model (see Table 1), suggesting that maternal effects did not affect the expression of those traits.

### How do population differences and maternal effects interact with ontogeny?

To determine how traits changed through offspring development, we compared each phenotype across age groups. ‘Age’ was a significant predictor of growth rate and burst swimming, indicating that these traits changed throughout ontogeny, while the other phenotypes were not affected by age. Across populations and food treatments, fry grew at a slower relative rate (*χ*^2^_1_ = 147.91; *P* < 0.001, CI: –0.010, –0.007; ***η***_p_^2^ = 0.50) and performed better in burst swimming trials (*χ*^2^_1_ = 30.178; *P* < 0.001; CI: 0.024, 0.049; ***η***_p_^2^ = 0.16) as they got older (Table 1). Burst swimming scores were also lower in sulfidic individuals (see above), but the interaction term ‘Population × Age’ was not significant, demonstrating that both populations exhibited similar changes in burst swimming throughout ontogeny.

To ensure that signals of population differentiation and maternal effects occurring at birth were not obscured by measurements later in life, we also subset our dataset for each phenotype to obtain only the earliest data point for each family. As mentioned above, fry from the sulfidic population and the low-food maternal treatment had significantly larger size at birth (Figure 2; Table 1). However, there were no population differences or maternal effects on brood size, which was higher in larger mothers (Mother SL; *F*_1,37_ = 23.342; *P* < 0.001; CI: 0.73, 1.79; ***η***_p_^2^ = 0.38) and mothers who spent longer in the food treatment (Treatment Length; *F*_1,37_ = 8.926; *P* = 0.005; CI: 0.07, 0.40; ***η***_p_^2^ = 0.19). Growth rate at birth was significantly lower in sulfidic fry (*F*_1,37_ = 5.025; *P* = 0.032; CI: –0.12, –0.006; ***η***_p_^2^ = 0.12; Table 1; Figure 3A), but there was no evidence for maternal effects on growth rate at birth (‘Food Treatment’ was not included in best-supported model). Note that the population difference in growth rate at birth disappeared as fry developed (Figure 3B). All other traits that were measured throughout ontogeny (burst swimming, exploratory behavior, and feeding accuracy) did not exhibit significant population differences or maternal effects at birth. Interestingly, the maternal effect that we detected on exploratory behavior across all ages (see above) was not evident at birth (*F*_1,31_ = 2.727; *P* = 0.108; CI: –0.29, 2.81). These results collectively demonstrate that—contrary to our hypothesis regarding ontogenetic variation of functional traits—population differentiation did not increase with age, and maternal effects were not always observable at birth, nor did they decline with age.

### How do population differences interact with maternal effects?

We hypothesized that variation from maternal effects would be aligned with population differences, but that there would be a significant interaction between population differences and maternal effects (*i.e*., different magnitudes of maternal effects in each population due to different evolutionary histories associated with resource stress). Contrary to our predictions, no ‘Population × Food Treatment’ interactions were included in the best-supported model for any of the traits we measured (Table 1), suggesting a general lack of support for significant interactions between population differences and maternal effects. Therefore, maternal effects—if present—were similar in direction and magnitude between populations.

To further address our hypothesis, we asked whether population differences and maternal effects were, in fact, aligned, and whether they explained a similar proportion of phenotypic variance when they acted in unison. We compared the signs (positive vs. negative coefficient estimates) and effect sizes (***η***_p_^2^) of population differences and maternal effects for the two traits with evidence of both effects simultaneously impacting trait expression—size at birth and exploratory behavior. For both traits, the difference between the nonsulfidic and sulfidic population occurred in the same direction as the trait shifts between the high- and low-food treatments, indicating that the effects were aligned (Figures 2A and 3E). Effect size estimates for ‘Population’ and ‘Food Treatment’ for both traits indicated that ‘Population’ had a larger effect on trait expression than ‘Food Treatment’ (***η***_p_^2^= 0.36 *vs*. 0.11 for size at birth and 0.28 vs. 0.18 for exploratory behavior).

### Multivariate analysis

In addition to the univariate analyses of trait variation, we were also interested in understanding how the traits covaried with one another, and whether and how multivariate phenotypes were impacted by population differences and maternal effects. We averaged each phenotype across ages for each family, conducted a PCA, and then analyzed principal component scores along the first two principal component axes. The first principal component accounted for 28.5% of variance in multivariate phenotypes, and scores along the first principal component were positively correlated with brood size, growth rate, and feeding accuracy, and they were negatively correlated with size at birth, age at maturity, burst swimming, and exploratory behavior (see Table S2C). The second principal component explained 21.1% of variance and was positively correlated with burst swimming and size at birth and negatively correlated with brood size, growth rate, feeding accuracy, exploratory behavior, and age at maturity (Table S2). After conducting model selection, the best-supported model for PC1 contained the main effects of ‘Population’, ‘Food Treatment’, and ‘Treatment Length’, and the best-supported model for PC2 contained the main effects of ‘Population’, ‘Mother SL’, and ‘Treatment Length’.

Sulfidic offspring had significantly lower scores along PC1 (*F*_1,24_ = 16.266; *P* < 0.001; CI: – 2.40, –0.78; ***η***_p_^2^ = 0.40), as did offspring from low-food mothers (*F*_1,24_ = 4.746; *P* = 0.039; CI: –1.59, –0.04), indicating population differences and maternal effects on PC1 scores. As for the univariate analyses, maternal effects were aligned with population differences in PC1 scores (Figure 4), ‘Population’ had a larger effect than ‘Food Treatment’ (***η***_p_^2^ for ‘Population’ = 0.40; ***η***_p_^2^ for ‘Food Treatment’ = 0.17), and there was no evidence for a significant ‘Population × Food Treatment’ interaction (the term was not included in best-supported model). We also found that sulfidic individuals exhibited marginally lower PC2 scores (*F*_1,24_ = 3.321; *P* = 0.081; CI: –1.45, 0.09; ***η***_p_^2^ = 0.12) but detected no evidence for maternal effects (the ‘Food Treatment’ term was not included in the best-supported model) on PC2.

**Figure 4.**
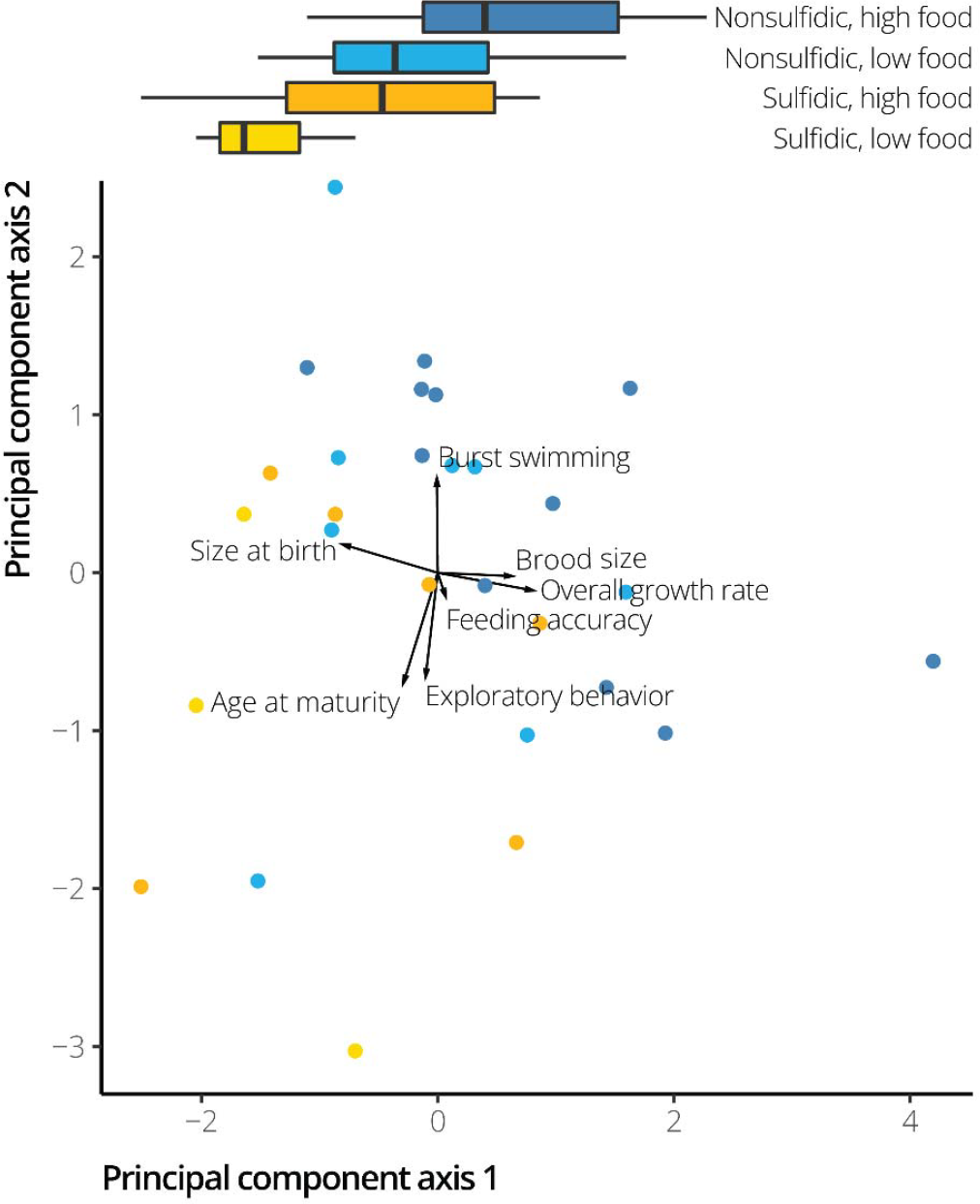
Plot of principal component scores representing linear combinations of all phenotypes in multivariate space. Scores were plotted along the first two principal components. Nonsulfidic families are shown in shades of blue, while sulfidic families are shown in shades of yellow. The high-food treatment within each population is shown in the darker shade (*i.e*., dark blue or dark yellow). Radiating from the origin are arrows that represent the correlation between the principal component scores and each input variable (shown as text at the end of each arrow). Loadings were calculated by multiplying the eigenvector for each input variable by the square-root of the eigenvalue for that principal component axis. At the top of the figure are boxplots that show the distribution of principal component scores along the first principal component axis, which exhibited significant effects of ‘Population’ and ‘Food Treatment’.

## Discussion

Phenotypic variation can be shaped by multiple genetic and nongenetic factors, but the interplay of genes and environmental effects is rarely disentangled in natural systems, even though it fundamentally impacts our inference of adaptation. We examined how genetic variation (*i.e*., population differences), phenotypic plasticity mediated by maternal effects, and their interactions shape trait expression in two populations of *Poecilia mexicana* that are exposed to strong divergent selection in nature. We found significant trait differences between populations even though fish were housed under common-garden conditions in the laboratory for at least three generations. In contrast, exposure of mothers to different food treatments impacted relatively few traits in their offspring and, if they occurred, had weaker effects than the population differences. In addition, maternal effects tended to be aligned with population differences and act in the same way across populations. Principal component analysis supported these overall conclusions in multivariate trait space. We also found no evidence for interactions between populations and food treatments, suggesting that while populations have diverged in phenotypic traits, they have retained similar maternal influences on those same traits. Overall, we found that the stark phenotypic differences between populations of *P. mexicana* that are evident in nature are largely a consequence of genetic divergence, likely representing local adaptation to the distinct ecological conditions of their habitats. Maternal effects in response to resource availability, while present in some traits, appear to accentuate population differences, but not cause them.

### Trait variation across populations and maternal food treatments

There is a rich history integrating field-based studies that quantify trait variation of poeciliid fishes in nature with laboratory-based studies that isolate causative environmental factors (Endler, 1980, 1995; Reznick & Bryga, 1987; Reznick, Bryga, & Endler, 1990; Langerhans, Gifford, & Joseph, 2007; Tobler *et al*., 2008; Ghalambor *et al*., 2015; Ingley & Johnson, 2016). While most studies focus on single or a few related traits and take a snapshot at a single ontogenetic stage, our study demonstrated that multiple complex trait differences quantified in common-garden conditions—across food treatments and across ontogeny—closely mirror trait differences between populations in nature.

First, our study corroborates a genetic basis for population divergence in reproductive life history traits. We documented significant population divergence in size at birth, with sulfidic mollies giving birth to larger offspring (see Figures 2A and 4). This finding is consistent with life history studies of sulfide spring populations in *P. mexicana* and other poeciliid species (Riesch *et al*., 2010b; Riesch, Plath, & Schlupp, 2010c; Riesch *et al*., 2014), suggesting that divergence in offspring size between sulfidic and nonsulfidic populations is a repeatable evolutionary consequence of sulfide spring colonization that is driven by natural selection for larger offspring size. Among-population variation in size at birth has also been documented in other poeciliids that are exposed to other sources of divergent selection, including variation in predation (Weeks & Gaggiotti, 1993; Reznick, Rodd, & Cardenas, 1996b; Schrader & Travis, 2012; Belk, Ingley, & Johnson, 2020). Our multivariate results (Figure 4) also indicated that, like other poeciliids, the nonsulfidic population of *P. mexicana* closely resembles an opportunistic life history strategy (Winemiller & Rose, 1992), which places a premium on earlier maturity and higher fecundity, at the expense of lower juvenile survivorship. The population differences we found in the sulfidic population may represent a shift in the direction of the equilibrium strategy, which is energetically costly to parents but should ultimately benefit offspring competitive ability. In natural populations, maternal effects induced by variation in resource availability likely accentuate genetic differences in offspring size across sulfidic and nonsulfidic habitats. This pattern was also evident across a plethora of life history traits in guppies; in all ten life history traits with evidence for significant genetic divergence and maternal effects, these effects always occurred in the same direction (Felmy *et al*., 2022).

Second, we found patterns of population differentiation in age at maturity that match with previous work in livebearing fishes. Prior studies have demonstrated that guppies in streams with high predation pressure on adults mature earlier than conspecifics from low-predation populations (Reznick & Endler, 1982). Several studies have also documented genetic differences in age at maturity between populations of *P. mexicana*. For example, cave mollies mature significantly later than surface populations, and there was also significant plasticity in age at maturity due to resource availability (Riesch *et al*., 2016a). Another study of surface mollies that compared size at maturity in sulfidic and nonsulfidic populations found that sulfidic individuals reached maturity at a significantly smaller size than nonsulfidic individuals (Riesch *et al*., 2011b), likely because energy limitation imposed by constraints associated with compensatory behaviors in sulfidic habitats (Tobler *et al*., 2009) selects for reduced growth rates and smaller size at maturity (Passow *et al*., 2017b). Since poeciliid males exhibit determinate growth that ends at maturity (Stearns & Sage, 1980), size at maturity should closely mirror age at maturity, assuming growth rates are the same as we found in our study. However, we found that sulfidic mollies matured at a significantly later age than nonsulfidic individuals, despite identical growth rates across ontogeny (Figure 3B). This discrepancy between our findings and those of previous studies suggests that plasticity in age at maturity may be strong and that variation in experimental design and rearing conditions matters.

Third, we found significant behavioral differences between populations from sulfidic and nonsulfidic habitats, including exploratory behavior, feeding accuracy, and burst swimming. Previous work has shown that nonsulfidic mollies—and ones that were better fed—were bolder in their natural habitat, but behavioral differences also disappeared in the laboratory (Riesch *et al*., 2009). While we did not measure boldness *per se*, exploratory behavior as measured in our experiment (*i.e*., activity levels in a novel arena) is often characterized as part of a behavioral syndrome that correlates with boldness (Conrad *et al*., 2011). Unlike previous work, our study found that individuals from the sulfidic population and the low-food treatment were more exploratory (Table 1 and Figure 3E). These results support that maternal resource environment affects exploratory behavior, but also imply heritable differences between populations. Similar heritable population differences were also found for feeding accuracy and burst swimming but without any evidence for maternal effects. At least for burst swimming, the pattern of population differentiation in our experiment again mirrors findings from adult fish in natural habitats (Camarillo *et al*., 2020).

### Trait variation and adaptive function

Sulfide springs and adjacent nonsulfidic habitats do not only differ in the presence and absence of H_2_S, but they also vary in numerous abiotic and biotic factors that are often unaddressed in studies of adaptation. Sulfidic habitats have lower dissolved oxygen concentrations, higher temperature, higher specific conductivity, and lower pH (Tobler *et al*., 2011), which in turn affect the biotic communities (Greenway *et al*., 2014) and selection associated with resource exploitation, competition, predation, and parasites (Riesch *et al*., 2010a; Tobler *et al*., 2014, 2015). Adaptation in sulfide springs is therefore not solely in response to selection from H_2_S but is instead in response to a multifarious selective regime that has caused multivariate phenotypic differentiation between populations (Tobler *et al*., 2018). The complexity of selective regimes and evolutionary responses make disentangling cause-and-effect relationships difficult, especially because theoretical predictions for the effects of different, covarying sources of selection are not mutually exclusive. For example, the evolution of large offspring size at birth could be explained by (1) selection from H_2_S, which should favor larger offspring with a lower surface-to-volume ratio to reduce the influx of toxic H_2_S (Riesch *et al*. 2014); (2) selection from resource constraints, which favors larger offspring with higher energy stores (Reznick *et al*., 1996a); or (3) relaxation of selection from predation, which also favors larger offspring (Reznick & Endler, 1982; Johnson & Belk, 2001; Jennions *et al*., 2006). Similarly, resource constraints and low predation also favor more exploratory individuals that are better able to locate and exploit resources under those conditions (Teska, Smith, & Novak, 1990; Kaun *et al*., 2007; Huang, Sieving, & Mary, 2012). Assessing the adaptive value of trait differences between sulfidic and nonsulfidic population is consequently nontrivial and remains a work in progress.

Variation in some traits investigated in our study may also be the consequence of genetic, developmental, or functional tradeoffs with other traits. Such tradeoffs are common in organisms inhabiting contrasting environments because divergent selection acting to optimize one trait can inadvertently influence other traits due to constraints (Ghalambor *et al*., 2004; Kawecki & Ebert, 2004; Garland, Downs, & Ives, 2022). For example, reductions in burst swimming performance—as documented in our study for fish from the sulfidic population—may arise as a consequence of selection for increased steady swimming efficiency, as different body shapes are associated with optimization of steady vs. unsteady swimming (Langerhans, 2007, 2009; Tokić & Yue, 2012). Indeed, a tradeoff between burst speed and sustained swimming performance has been documented in adult fish from the same populations we studied here. The tradeoff is likely balanced by the need for energy-efficient swimming in sulfidic habitats with resource constraints (but low predation) and selection for efficient predator avoidance in nonsulfidic habitats with high abundances of natural enemies (but abundant resources) (Camarillo *et al*., 2020). Our study found a similar reduction in burst swimming performance in the sulfidic population, even among individuals that had not yet reached maturity, suggesting population differences in burst swimming arise early in ontogeny. Our results also matched *a priori* predictions regarding tradeoffs between respiration and feeding. Habitats rich in H_2_S also experience rampant hypoxia, which has selected for the evolution of craniofacial traits (larger heads and jaws and longer gill filaments) that increase ventilation efficiency (Camarillo *et al*., 2020). In addition to changes in morphology, sulfidic individuals also exhibit decreased foraging efficiency compared to nonsulfidic individuals (Tobler *et al*., 2009), which was supported by our findings related to feeding accuracy (Figure 3F). The decreases in feeding efficiency noted in this and other studies may therefore be a consequence of the craniofacial modifications that accompany colonization of sulfidic habitats.

### Plasticity accentuates genetic trait differentiation in natural populations

Phenotypes in nature are the sum of genetic and environmental effects, but, surprisingly, we found no evidence for canalization or the evolution of plasticity by genotype × environment interactions. Our work demonstrated that maternal effects were aligned with population differences, accentuating trait divergence between populations, and that the trait shifts induced by maternal effects were of a similar magnitude in both populations. For both traits in which we observed significant population differentiation and maternal effects (size at birth and exploratory behavior), the lowest phenotypic scores were found in nonsulfidic fry from high-food mothers, and the highest scores were found in sulfidic fry from low-food mothers (Figures 2A and 3E). Because sulfidic mollies exhibit reduced foraging efficiency and body condition as a consequence of hypoxia (Tobler, 2008; Tobler *et al*., 2009), sulfidic habitats are naturally analogous to our low-food treatment, and nonsulfidic habitats are similar to the high-food treatment. If maternal effects enhance population differences in natural populations similarly as in our experiments, then it could explain why stronger trait divergence is typically observed in nature than in common garden reared fish (Tobler *et al*., 2008; Passow *et al*., 2015). Likewise, it is important to note that our experiments only captured a small aspect of plasticity—the portion controlled by the mother. Future work needs to address genetic effects, maternal effects, and plasticity in response to variation in environmental factors directly experienced by the offspring to better understand the forces driving trait variation in nature. We also caution that the shifts found in our study are only from comparisons of a single population pair. Our understanding of the relative roles of genetic and plastic effects on phenotypic variation would benefit greatly from studies that expand to other sulfidic and nonsulfidic population pairs in *P. mexicana* and other species.

## Acknowledgements

We would like to thank the members of the community in Tapijulapa, Tabasco for access to field sites and for their hospitality. We also would like to thank the Mexican government for issuing permits for past field collections, as well as our collaborators at the Universidad Juárez Autónoma de Tabasco and the Centro de Investigación E Innovación Para la Enseñanza y El Aprendizaje for support of research expeditions to obtain the specimens that gave rise to the populations used in this study. The assistance provided by J. Onnen and Q. La Fon was essential for processing our diverse datasets. Funding for this research was provided by grants from the National Science Foundation (IOS-1931657), the U.S. Army Research Office (W911NF-15-1-0175), and the Des Lee Collaborative Vision in Zoological Studies to MT, and by a Graduate Assistance in Areas of National Need (GAANN) fellowship from the U.S. Department of Education to JLC.

## Data Availability Statement

All datasets and R scripts used to analyze the data, with associated documentation, have been archived on GitHub (https://github.com/michitobler/common-garden).

