## Supplementary material for "Genetics and resource availability shape divergence in life history and behavior between locally-adapted populations of Atlantic mollies (*Poecilia mexicana*, Poeciliidae)"

Table S1. Model terms in the best-supported model for each dependent variable using all broods, or only broods where the female was in the food treatment for at least seven days. Empty cells represent terms that were not present in the best-supported model for that dependent variable. Type-III ANOVA or Wald’s Chi-square tests were used to determine whether model terms were significantly different from zero. *F*-statistics or *c*^2^ statistics are presented for each included model term, with the degrees of freedom in a subscript. *P*-values are presented underneath each test statistic. Note that for the models using a seven-day cutoff, the treatment length was not included in the AIC.

| **Dependent Variable** | **Brood Size** | **Treatment Length** | **Mother SL** | **Age** | **Population** | **Food Treatment** |
| --- | --- | --- | --- | --- | --- | --- |
| Size at Birth (All) | *F*_1,35_ = 20.876  *P* < 0.001 |  | *F*_1,35_ = 16.593  *P* < 0.001 |  | *F*_1,35_ = 20.032  *P* < 0.001 | *F*_1,35_ = 4.354  *P* = 0.044 |
| Size at Birth (Seven Day) | *F*_1,25_ = 15.583  *P* < 0.001 |  | *F*_1,25_ = 5.873  *P* < 0.023 |  | *F*_1,25_ = 16.025  *P* < 0.001 | *F*_1,25_ = 2.859  *P* = 0.103 |
| Brood Size (All) |  | *F*_1,37_ = 8.926  *P =* 0.005 | *F*_1,37_ = 23.342  *P* < 0.001 |  |  |  |
| Brood Size (Seven Day) |  |  | *F*_1,28_ = 10.376  *P* < 0.003 |  |  |  |
| Age at Maturity (All) |  |  |  |  | *F*_1,26_ = 4.420  *P =* 0.045 |  |
| Age at Maturity (Seven Day) |  |  |  |  |  | *F*_1,18_ = 0.828  *P =* 0.375 |
| Growth Rate at Birth (All) |  |  |  |  | *F*_1,37_ = 5.025  *P =* 0.031 |  |
| Growth Rate at Birth (Seven Day) |  |  |  |  | *F*_1,28_ = 3.127  *P =* 0.088 |  |
| Overall Growth Rate (All) |  |  |  | χ^2^_1_ = 147.91  *P* < 0.001 |  |  |
| Overall Growth Rate (Seven Day) |  |  |  | χ^2^_1_ = 113.75  *P* < 0.001 |  |  |
| Burst Swimming at Birth (All) |  |  | *F*_1,39_ = 5.025  *P =* 0.121 |  |  |  |
| Burst Swimming at Birth (Seven Day) |  |  |  |  | *F*_1,28_ = 0.910  *P =* 0.348 |  |
| Overall Burst Swimming (All) |  |  |  | χ^2^_1_ = 30.178  *P* < 0.001 | χ^2^_1_ = 4.308  *P* = 0.038 |  |
| Overall Burst Swimming (Seven Day) |  |  |  | χ^2^_1_ = 29.498  *P* < 0.001 | χ^2^_1_ = 3.812  *P* = 0.051 |  |

Table S2: Descriptive information for principal component (PC) analyses for multiple traits. Loadings and eigenvalues for each PC axis for each multivariate phenotype. The cumulative proportion of variance explained by each PC axis is given at the base of each column for each phenotype. Loadings were calculated as the eigenvector multiplied by the square root of the eigenvalue for each PC axis and represent the correlation coefficient between the original phenotypic data and the linearly transformed principal component scores. The table contains information for PC analyses that were conducted for (A) Burst Swimming, (B) Exploratory Behavior, and (C) Multivariate Trait Variation.

| **Phenotype** | **Input Variables** | **PC1** | **PC2** | **PC3** | **PC4** | **PC5** | **PC6** | **PC7** |
| --- | --- | --- | --- | --- | --- | --- | --- | --- |
| **(A) Burst swimming** | *V*_max_ | 0.963 | -0.029 | -0.270 |  |  |  |  |
|  | *A*_max_ | 0.911 | -0.380 | 0.159 |  |  |  |  |
|  | *D*_net_ | 0.901 | 0.416 | 0.127 |  |  |  |  |
|  | Eigenvalue | 2.570 | 0.318 | 0.114 |  |  |  |  |
|  | Variance Explained | 0.856 | 0.962 | 1.000 |  |  |  |  |
| **(B) Exploratory Behavior** | *V*_avg_ | 0.833 | 0.469 | -0.252 | -0.151 | 0.006 |  |  |
|  | *V*_max_ | 0.809 | -0.581 | 0.041 | 0.015 | 0.073 |  |  |
|  | *A*_max_ | 0.804 | -0.587 | 0.055 | -0.012 | -0.073 |  |  |
|  | *D*_total_ | 0.819 | 0.486 | -0.265 | 0.149 | -0.007 |  |  |
|  | *D*_center_ | 0.420 | 0.365 | 0.831 | 0.002 | 0.001 |  |  |
|  | Eigenvalue | 2.842 | 1.273 | 0.829 | 0.045 | 0.011 |  |  |
|  | Variance Explained | 0.569 | 0.823 | 0.989 | 0.998 | 1.000 |  |  |
| **(C) Multivariate Trait Variation** | Size at Birth | -0.844 | 0.188 | -0.167 | -0.111 | -0.365 | -0.181 | 0.215 |
|  | Brood Size | 0.672 | -0.023 | -0.304 | 0.300 | -0.174 | -0.578 | 0.011 |
|  | Age at Maturity | -0.302 | -0.728 | -0.096 | 0.129 | 0.562 | -0.151 | 0.121 |
|  | Growth Rate | 0.850 | -0.118 | -0.006 | -0.418 | -0.038 | 0.194 | 0.225 |
|  | Burst Swimming | -0.007 | 0.628 | 0.430 | -0.374 | 0.390 | -0.357 | 0.015 |
|  | Exploratory Behavior | -0.106 | -0.688 | 0.240 | -0.563 | -0.276 | -0.226 | -0.116 |
|  | Feeding Accuracy | 0.080 | -0.178 | 0.847 | 0.448 | -0.191 | -0.014 | 0.087 |
|  | Eigenvalue | 1.995 | 1.480 | 1.090 | 0.951 | 0.746 | 0.607 | 0.133 |
|  | Variance Explained | 0.285 | 0.496 | 0.652 | 0.788 | 0.894 | 0.981 | 1.000 |

Table S3. Full model selection tables for each phenotype measured. Models were weighted by AICc, and the best-supported model (with a delta value of 0) was used in analyses. Continuous variables that are included in models are denoted with the regression coefficient for that model term, and categorical variables that were included are denoted with a '+'. Empty cells represent terms that were not present in the model.

| **Dependent Variable** | **Intercept** | **Brood Size** | **Treatment Length** | **Mother SL** | **Age** | **Population** | **Food Treatment** | **Age × Population** | **Age × Food Treatment** | **Population × Food Treatment** | **Age × Population × Food Treatment** | **df** | **logLik** | **AICc** | **delta** | **weight** |
| --- | --- | --- | --- | --- | --- | --- | --- | --- | --- | --- | --- | --- | --- | --- | --- | --- |
| *Birth Size* | 6.910 | -0.033 |  | 0.059 |  | + | + |  |  |  |  | 6 | -32.2 | 78.9 | 0.0 | 0.377 |
|  | 8.161 | -0.030 | -0.009 | 0.040 |  | + |  |  |  |  |  | 6 | -32.3 | 79.1 | 0.3 | 0.333 |
|  | 7.413 | -0.037 |  | 0.055 |  | + |  |  |  |  |  | 5 | -34.5 | 80.8 | 1.9 | 0.145 |
|  | 9.948 | -0.018 | -0.014 |  |  | + |  |  |  |  |  | 5 | -35.5 | 82.8 | 3.9 | 0.054 |
|  | 9.727 | -0.014 | -0.014 |  |  | + | + |  |  |  |  | 6 | -34.2 | 83.0 | 4.1 | 0.048 |
|  | 9.388 |  | -0.016 |  |  | + | + |  |  |  |  | 5 | -36.5 | 84.8 | 6.0 | 0.019 |
|  | 8.601 |  | -0.014 | 0.015 |  | + | + |  |  |  |  | 6 | -36.0 | 86.5 | 7.6 | 0.008 |
|  | 9.335 |  | -0.015 |  |  | + | + |  |  | + |  | 6 | -36.3 | 87.2 | 8.4 | 0.006 |
|  | 9.615 |  | -0.016 |  |  | + |  |  |  |  |  | 4 | -39.3 | 87.7 | 8.8 | 0.005 |
|  | 9.485 |  | -0.016 | 0.003 |  | + |  |  |  |  |  | 5 | -39.3 | 90.3 | 11.4 | 0.001 |
|  | 9.771 | -0.023 |  |  |  | + |  |  |  |  |  | 4 | -40.7 | 90.6 | 11.8 | 0.001 |
|  | 9.565 | -0.020 |  |  |  | + | + |  |  |  |  | 5 | -39.9 | 91.6 | 12.7 | 0.001 |
|  | 8.676 | -0.034 |  | 0.034 |  |  |  |  |  |  |  | 4 | -41.7 | 92.5 | 13.6 | 0.000 |
|  | 10.244 | -0.023 | -0.008 |  |  |  |  |  |  |  |  | 4 | -42.1 | 93.3 | 14.5 | 0.000 |
|  | 9.484 | -0.020 |  |  |  | + | + |  |  | + |  | 6 | -39.4 | 93.4 | 14.5 | 0.000 |
|  | 8.483 | -0.032 |  | 0.035 |  |  | + |  |  |  |  | 5 | -41.2 | 94.2 | 15.3 | 0.000 |
|  | 9.159 | -0.030 | -0.005 | 0.025 |  |  |  |  |  |  |  | 5 | -41.2 | 94.2 | 15.3 | 0.000 |
|  | 10.061 | -0.025 |  |  |  |  |  |  |  |  |  | 3 | -43.8 | 94.3 | 15.4 | 0.000 |
|  | 7.552 |  |  | 0.029 |  | + | + |  |  |  |  | 5 | -41.5 | 94.8 | 15.9 | 0.000 |
|  | 10.136 | -0.021 | -0.008 |  |  |  | + |  |  |  |  | 5 | -41.8 | 95.4 | 16.6 | 0.000 |
|  | 9.042 |  |  |  |  | + | + |  |  |  |  | 4 | -43.3 | 95.7 | 16.8 | 0.000 |
|  | 8.944 | -0.029 | -0.004 | 0.027 |  |  | + |  |  |  |  | 6 | -40.8 | 96.1 | 17.3 | 0.000 |
|  | 9.948 | -0.023 |  |  |  |  | + |  |  |  |  | 4 | -43.5 | 96.2 | 17.4 | 0.000 |
|  | 7.532 |  |  | 0.029 |  | + | + |  |  | + |  | 6 | -41.2 | 96.9 | 18.1 | 0.000 |
|  | 8.961 |  |  |  |  | + | + |  |  | + |  | 5 | -42.9 | 97.5 | 18.6 | 0.000 |
|  | 9.276 |  |  |  |  | + |  |  |  |  |  | 3 | -45.6 | 97.8 | 19.0 | 0.000 |
|  | 8.490 |  |  | 0.016 |  | + |  |  |  |  |  | 4 | -45.1 | 99.3 | 20.4 | 0.000 |
|  | 9.863 |  | -0.011 |  |  |  |  |  |  |  |  | 3 | -46.4 | 99.5 | 20.7 | 0.000 |
|  | 9.692 |  | -0.010 |  |  |  | + |  |  |  |  | 4 | -45.2 | 99.6 | 20.7 | 0.000 |
|  | 10.520 |  | -0.012 | -0.013 |  |  |  |  |  |  |  | 4 | -46.1 | 101.3 | 22.5 | 0.000 |
|  | 9.390 |  |  |  |  |  | + |  |  |  |  | 3 | -47.4 | 101.5 | 22.7 | 0.000 |
|  | 10.033 |  | -0.011 | -0.007 |  |  | + |  |  |  |  | 5 | -45.1 | 102.0 | 23.2 | 0.000 |
|  | 9.067 |  |  | 0.007 |  |  | + |  |  |  |  | 4 | -47.4 | 103.8 | 25.0 | 0.000 |
|  | 9.562 |  |  | 0.000 |  |  |  |  |  |  |  | 3 | -48.8 | 104.3 | 25.4 | 0.000 |
| *Brood Size* | -44.963 |  | 0.240 | 1.263 |  |  |  |  |  |  |  | 4 | -155.4 | 320.0 | 0.0 | 0.461 |
|  | -38.057 |  | 0.222 | 1.168 |  |  | + |  |  |  |  | 5 | -154.6 | 321.0 | 1.0 | 0.279 |
|  | -44.511 |  | 0.242 | 1.256 |  | + |  |  |  |  |  | 5 | -155.4 | 322.6 | 2.6 | 0.125 |
|  | -35.987 |  | 0.228 | 1.136 |  | + | + |  |  |  |  | 6 | -154.5 | 323.6 | 3.7 | 0.074 |
|  | -18.203 |  |  | 0.890 |  |  | + |  |  |  |  | 4 | -158.4 | 325.9 | 5.9 | 0.024 |
|  | -25.918 |  |  | 0.996 |  |  |  |  |  |  |  | 3 | -159.7 | 326.2 | 6.2 | 0.021 |
|  | -29.434 |  |  | 1.050 |  | + |  |  |  |  |  | 4 | -159.6 | 328.3 | 8.4 | 0.007 |
|  | -19.501 |  |  | 0.909 |  | + | + |  |  |  |  | 5 | -158.4 | 328.5 | 8.5 | 0.007 |
|  | -19.401 |  |  | 0.914 |  | + | + |  |  | + |  | 6 | -158.3 | 331.2 | 11.3 | 0.002 |
|  | 24.042 |  |  |  |  |  | + |  |  |  |  | 3 | -163.4 | 333.6 | 13.6 | 0.001 |
|  | 26.478 |  |  |  |  | + | + |  |  |  |  | 4 | -162.9 | 334.9 | 14.9 | 0.000 |
|  | 21.308 |  | 0.091 |  |  |  | + |  |  |  |  | 4 | -162.9 | 335.0 | 15.0 | 0.000 |
|  | 23.551 |  | 0.133 |  |  | + | + |  |  |  |  | 5 | -161.8 | 335.4 | 15.4 | 0.000 |
|  | 16.788 |  | 0.108 |  |  |  |  |  |  |  |  | 3 | -165.2 | 337.1 | 17.1 | 0.000 |
|  | 26.417 |  |  |  |  | + | + |  |  | + |  | 5 | -162.9 | 337.5 | 17.5 | 0.000 |
|  | 21.409 |  |  |  |  | + |  |  |  |  |  | 3 | -165.6 | 337.9 | 17.9 | 0.000 |
|  | 23.156 |  | 0.135 |  |  | + | + |  |  | + |  | 6 | -161.8 | 338.1 | 18.2 | 0.000 |
|  | 18.408 |  | 0.144 |  |  | + |  |  |  |  |  | 4 | -164.5 | 338.2 | 18.2 | 0.000 |
| *Growth Rate* | -2.829 |  |  |  | -0.020 |  |  |  |  |  |  | 4 | -106.0 | 220.2 | 0.0 | 0.856 |
|  | -2.818 |  |  |  | -0.020 |  | + |  |  |  |  | 5 | -107.5 | 225.4 | 5.2 | 0.062 |
|  | -2.824 |  |  |  | -0.020 | + |  |  |  |  |  | 5 | -107.5 | 225.5 | 5.3 | 0.061 |
|  | -2.907 | 0.004 |  |  | -0.020 |  |  |  |  |  |  | 5 | -109.9 | 230.2 | 10.0 | 0.006 |
|  | -2.648 |  |  | -0.004 | -0.021 |  |  |  |  |  |  | 5 | -110.1 | 230.6 | 10.4 | 0.005 |
|  | -2.810 |  |  |  | -0.020 | + | + |  |  |  |  | 6 | -109.0 | 230.7 | 10.5 | 0.004 |
|  | -2.862 |  | 0.002 |  | -0.021 |  |  |  |  |  |  | 5 | -110.7 | 231.9 | 11.7 | 0.003 |
|  | -2.819 |  |  |  | -0.020 | + | + |  |  | + |  | 7 | -109.9 | 234.7 | 14.5 | 0.001 |
|  | -2.914 | 0.004 |  |  | -0.020 |  | + |  |  |  |  | 6 | -111.4 | 235.4 | 15.2 | 0.000 |
|  | -2.901 | 0.004 |  |  | -0.020 | + |  |  |  |  |  | 6 | -111.4 | 235.5 | 15.3 | 0.000 |
|  | -2.582 |  |  | -0.005 | -0.020 | + |  |  |  |  |  | 6 | -111.5 | 235.5 | 15.3 | 0.000 |
|  | -2.605 |  |  | -0.004 | -0.020 |  | + |  |  |  |  | 6 | -111.5 | 235.7 | 15.5 | 0.000 |
|  | -2.849 |  | 0.003 |  | -0.021 | + |  |  |  |  |  | 6 | -111.7 | 236.0 | 15.8 | 0.000 |
|  | -2.778 |  |  |  | -0.022 | + |  | + |  |  |  | 6 | -111.9 | 236.4 | 16.2 | 0.000 |
|  | -2.857 |  |  |  | -0.019 |  | + |  | + |  |  | 6 | -111.9 | 236.5 | 16.3 | 0.000 |
|  | -2.855 |  | 0.002 |  | -0.021 |  | + |  |  |  |  | 6 | -112.3 | 237.2 | 17.0 | 0.000 |
|  | -2.476 | 0.007 |  | -0.010 | -0.021 |  |  |  |  |  |  | 6 | -112.5 | 237.7 | 17.5 | 0.000 |
|  | -2.500 |  |  | -0.006 | -0.020 | + | + |  |  |  |  | 7 | -112.8 | 240.3 | 20.1 | 0.000 |
|  | -2.907 | 0.004 |  |  | -0.020 | + | + |  |  |  |  | 7 | -113.0 | 240.7 | 20.5 | 0.000 |
|  | -2.839 |  | 0.003 |  | -0.020 | + | + |  |  |  |  | 7 | -113.3 | 241.4 | 21.2 | 0.000 |
|  | -2.304 | 0.008 |  | -0.013 | -0.021 | + |  |  |  |  |  | 7 | -113.3 | 241.5 | 21.3 | 0.000 |
|  | -2.849 |  |  |  | -0.019 | + | + |  | + |  |  | 7 | -113.5 | 241.7 | 21.5 | 0.000 |
|  | -2.770 |  |  |  | -0.022 | + | + | + |  |  |  | 7 | -113.5 | 241.7 | 21.5 | 0.000 |
|  | -2.929 | 0.004 | 0.002 |  | -0.021 |  |  |  |  |  |  | 6 | -114.8 | 242.2 | 22.0 | 0.000 |
|  | -2.794 |  | 0.002 | -0.001 | -0.021 |  |  |  |  |  |  | 6 | -115.0 | 242.6 | 22.4 | 0.000 |
|  | -2.480 | 0.007 |  | -0.010 | -0.021 |  | + |  |  |  |  | 7 | -114.1 | 243.0 | 22.8 | 0.000 |
|  | -2.857 | 0.004 |  |  | -0.022 | + |  | + |  |  |  | 7 | -115.9 | 246.5 | 26.3 | 0.000 |
|  | -2.914 | 0.004 | 0.002 |  | -0.020 | + |  |  |  |  |  | 7 | -115.9 | 246.5 | 26.3 | 0.000 |
|  | -2.951 | 0.004 |  |  | -0.019 |  | + |  | + |  |  | 7 | -115.9 | 246.5 | 26.3 | 0.000 |
|  | -2.548 |  |  | -0.005 | -0.022 | + |  | + |  |  |  | 7 | -115.9 | 246.5 | 26.3 | 0.000 |
|  | -2.717 |  | 0.002 | -0.003 | -0.021 | + |  |  |  |  |  | 7 | -115.9 | 246.6 | 26.4 | 0.000 |
|  | -2.642 |  |  | -0.004 | -0.019 |  | + |  | + |  |  | 7 | -116.0 | 246.7 | 26.5 | 0.000 |
|  | -2.809 |  | 0.003 |  | -0.022 | + |  | + |  |  |  | 7 | -116.2 | 247.2 | 27.0 | 0.000 |
|  | -2.943 | 0.004 | 0.002 |  | -0.021 |  | + |  |  |  |  | 7 | -116.3 | 247.5 | 27.3 | 0.000 |
|  | -2.761 |  | 0.002 | -0.002 | -0.021 |  | + |  |  |  |  | 7 | -116.5 | 247.9 | 27.7 | 0.000 |
|  | -2.890 |  | 0.002 |  | -0.019 |  | + |  | + |  |  | 7 | -116.8 | 248.4 | 28.2 | 0.000 |
|  | -2.504 | 0.007 | 0.000 | -0.009 | -0.021 |  |  |  |  |  |  | 7 | -117.9 | 250.6 | 30.4 | 0.000 |
|  | -3.340 |  |  |  |  | + |  |  |  |  |  | 4 | -150.4 | 309.0 | 88.8 | 0.000 |
|  | -3.251 |  |  |  |  | + | + |  |  |  |  | 5 | -150.5 | 311.3 | 91.1 | 0.000 |
|  | -3.375 |  |  |  |  |  | + |  |  |  |  | 4 | -151.7 | 311.7 | 91.5 | 0.000 |
|  | -3.229 |  |  |  |  | + | + |  |  | + |  | 6 | -150.9 | 314.4 | 94.2 | 0.000 |
|  | -4.008 |  |  | 0.012 |  |  |  |  |  |  |  | 4 | -153.9 | 316.0 | 95.8 | 0.000 |
|  | -3.564 | 0.007 |  |  |  |  |  |  |  |  |  | 4 | -154.6 | 317.5 | 97.3 | 0.000 |
|  | -3.458 | 0.007 |  |  |  | + |  |  |  |  |  | 5 | -153.7 | 317.9 | 97.7 | 0.000 |
|  | -3.718 |  |  | 0.008 |  | + |  |  |  |  |  | 5 | -153.9 | 318.1 | 97.9 | 0.000 |
|  | -3.886 |  |  | 0.011 |  |  | + |  |  |  |  | 5 | -154.7 | 319.8 | 99.6 | 0.000 |
|  | -3.377 |  | -0.003 |  |  |  |  |  |  |  |  | 4 | -155.8 | 319.8 | 99.6 | 0.000 |
|  | -3.481 |  |  | 0.004 |  | + | + |  |  |  |  | 6 | -154.2 | 321.1 | 100.9 | 0.000 |
|  | -3.331 |  | -0.001 |  |  | + |  |  |  |  |  | 5 | -155.4 | 321.3 | 101.1 | 0.000 |
|  | -3.361 | 0.005 |  |  |  | + | + |  |  |  |  | 6 | -154.4 | 321.3 | 101.1 | 0.000 |
|  | -3.498 | 0.006 |  |  |  |  | + |  |  |  |  | 5 | -155.4 | 321.3 | 101.1 | 0.000 |
|  | -3.297 |  | -0.003 |  |  |  | + |  |  |  |  | 5 | -156.1 | 322.6 | 102.4 | 0.000 |
|  | -3.239 |  | -0.001 |  |  | + | + |  |  |  |  | 6 | -155.5 | 323.6 | 103.4 | 0.000 |
|  | -3.459 |  |  | 0.004 |  | + | + |  |  | + |  | 7 | -154.7 | 324.2 | 104.0 | 0.000 |
|  | -3.340 | 0.005 |  |  |  | + | + |  |  | + |  | 7 | -154.8 | 324.3 | 104.1 | 0.000 |
|  | -3.913 | 0.004 |  | 0.009 |  |  |  |  |  |  |  | 5 | -158.0 | 326.3 | 106.1 | 0.000 |
|  | -3.218 |  | -0.001 |  |  | + | + |  |  | + |  | 7 | -156.0 | 326.7 | 106.5 | 0.000 |
|  | -3.516 | 0.006 |  | 0.001 |  | + |  |  |  |  |  | 6 | -157.5 | 327.6 | 107.4 | 0.000 |
|  | -3.894 |  | -0.001 | 0.010 |  |  |  |  |  |  |  | 5 | -158.9 | 328.1 | 107.9 | 0.000 |
|  | -3.498 | 0.007 | -0.003 |  |  |  |  |  |  |  |  | 5 | -158.9 | 328.3 | 108.1 | 0.000 |
|  | -3.447 | 0.007 | -0.001 |  |  | + |  |  |  |  |  | 6 | -158.7 | 330.1 | 109.9 | 0.000 |
|  | -3.830 | 0.003 |  | 0.008 |  |  | + |  |  |  |  | 6 | -158.9 | 330.3 | 110.1 | 0.000 |
|  | -3.725 |  | 0.000 | 0.008 |  | + |  |  |  |  |  | 6 | -158.9 | 330.4 | 110.2 | 0.000 |
|  | -3.355 | 0.005 |  | 0.000 |  | + | + |  |  |  |  | 7 | -158.1 | 331.1 | 110.9 | 0.000 |
|  | -3.698 |  | -0.002 | 0.008 |  |  | + |  |  |  |  | 6 | -159.5 | 331.7 | 111.5 | 0.000 |
|  | -3.422 | 0.006 | -0.003 |  |  |  | + |  |  |  |  | 6 | -159.7 | 332.0 | 111.8 | 0.000 |
|  | -3.453 |  | 0.000 | 0.004 |  | + | + |  |  |  |  | 7 | -159.3 | 333.3 | 113.1 | 0.000 |
|  | -3.350 | 0.005 | -0.001 |  |  | + | + |  |  |  |  | 7 | -159.4 | 333.5 | 113.3 | 0.000 |
|  | -3.637 | 0.006 | -0.002 | 0.003 |  |  |  |  |  |  |  | 6 | -162.5 | 337.7 | 117.5 | 0.000 |
|  | -3.419 | 0.007 | -0.001 | -0.001 |  | + |  |  |  |  |  | 7 | -162.4 | 339.6 | 119.4 | 0.000 |
|  | -3.508 | 0.005 | -0.003 | 0.002 |  |  | + |  |  |  |  | 7 | -163.4 | 341.5 | 121.3 | 0.000 |
| *Age at Maturity* | 43.684 |  |  |  |  | + |  |  |  |  |  | 3 | -109.2 | 225.3 | 0.0 | 0.155 |
|  | 63.776 |  |  | -0.413 |  | + |  |  |  |  |  | 4 | -108.1 | 225.9 | 0.5 | 0.118 |
|  | 72.625 |  |  | -0.542 |  |  |  |  |  |  |  | 3 | -109.6 | 226.2 | 0.8 | 0.101 |
|  | 41.364 |  | 0.113 |  |  | + |  |  |  |  |  | 4 | -108.6 | 227.0 | 1.7 | 0.067 |
|  | 45.438 | -0.077 |  |  |  | + |  |  |  |  |  | 4 | -109.0 | 227.7 | 2.4 | 0.046 |
|  | 44.074 |  |  |  |  | + | + |  |  |  |  | 4 | -109.1 | 228.0 | 2.7 | 0.040 |
|  | 77.613 |  |  | -0.611 |  |  | + |  |  |  |  | 4 | -109.2 | 228.1 | 2.7 | 0.039 |
|  | 43.726 |  | 0.148 |  |  |  |  |  |  |  |  | 3 | -110.6 | 228.1 | 2.8 | 0.038 |
|  | 67.329 |  | 0.093 | -0.474 |  |  |  |  |  |  |  | 4 | -109.3 | 228.3 | 2.9 | 0.036 |
|  | 59.700 |  | 0.077 | -0.362 |  | + |  |  |  |  |  | 5 | -107.8 | 228.4 | 3.0 | 0.034 |
|  | 67.903 |  |  | -0.471 |  | + | + |  |  |  |  | 5 | -107.8 | 228.4 | 3.1 | 0.033 |
|  | 44.173 | -0.190 | 0.188 |  |  | + |  |  |  |  |  | 5 | -107.8 | 228.4 | 3.1 | 0.033 |
|  | 46.925 | -0.238 | 0.236 |  |  |  |  |  |  |  |  | 4 | -109.4 | 228.6 | 3.3 | 0.030 |
|  | 74.005 | 0.052 |  | -0.595 |  |  |  |  |  |  |  | 4 | -109.5 | 228.8 | 3.5 | 0.027 |
|  | 64.743 | 0.034 |  | -0.449 |  | + |  |  |  |  |  | 5 | -108.0 | 228.8 | 3.5 | 0.027 |
|  | 49.262 | -0.100 |  |  |  |  |  |  |  |  |  | 3 | -111.1 | 229.3 | 3.9 | 0.022 |
|  | 47.765 |  |  |  |  |  | + |  |  |  |  | 3 | -111.3 | 229.6 | 4.3 | 0.018 |
|  | 41.585 |  | 0.112 |  |  | + | + |  |  |  |  | 5 | -108.6 | 230.0 | 4.6 | 0.015 |
|  | 46.957 | -0.102 |  |  |  | + | + |  |  |  |  | 5 | -108.9 | 230.5 | 5.2 | 0.011 |
|  | 72.649 |  | 0.075 | -0.547 |  |  | + |  |  |  |  | 5 | -109.0 | 230.6 | 5.3 | 0.011 |
|  | 44.223 |  | 0.145 |  |  |  | + |  |  |  |  | 4 | -110.5 | 230.8 | 5.5 | 0.010 |
|  | 49.558 | -0.290 | 0.244 |  |  |  | + |  |  |  |  | 5 | -109.1 | 230.9 | 5.6 | 0.010 |
|  | 43.636 |  |  |  |  | + | + |  |  | + |  | 5 | -109.1 | 230.9 | 5.6 | 0.010 |
|  | 77.787 | 0.010 |  | -0.620 |  |  | + |  |  |  |  | 5 | -109.2 | 231.0 | 5.7 | 0.009 |
|  | 61.533 | -0.102 | 0.147 | -0.330 |  |  |  |  |  |  |  | 5 | -109.2 | 231.1 | 5.7 | 0.009 |
|  | 46.211 | -0.230 | 0.196 |  |  | + | + |  |  |  |  | 6 | -107.6 | 231.3 | 5.9 | 0.008 |
|  | 63.786 |  | 0.065 | -0.418 |  | + | + |  |  |  |  | 6 | -107.7 | 231.3 | 6.0 | 0.008 |
|  | 67.965 |  |  | -0.484 |  | + | + |  |  | + |  | 6 | -107.7 | 231.4 | 6.1 | 0.007 |
|  | 53.778 | -0.104 | 0.132 | -0.215 |  | + |  |  |  |  |  | 6 | -107.7 | 231.4 | 6.1 | 0.007 |
|  | 51.483 | -0.139 |  |  |  |  | + |  |  |  |  | 4 | -110.9 | 231.6 | 6.2 | 0.007 |
|  | 67.985 | 0.005 |  | -0.475 |  | + | + |  |  |  |  | 6 | -107.8 | 231.7 | 6.3 | 0.007 |
|  | 41.311 |  | 0.110 |  |  | + | + |  |  | + |  | 6 | -108.6 | 233.2 | 7.8 | 0.003 |
|  | 65.082 | -0.148 | 0.151 | -0.349 |  |  | + |  |  |  |  | 6 | -108.8 | 233.5 | 8.2 | 0.003 |
|  | 46.608 | -0.109 |  |  |  | + | + |  |  | + |  | 6 | -108.8 | 233.6 | 8.3 | 0.002 |
| *Burst swimming* | -0.755 |  |  |  | 0.036 | + |  |  |  |  |  | 5 | -299.2 | 608.8 | 0.0 | 0.293 |
|  | -1.258 |  |  |  | 0.055 | + |  | + |  |  |  | 6 | -298.2 | 608.9 | 0.1 | 0.286 |
|  | -0.964 |  |  |  | 0.037 |  |  |  |  |  |  | 4 | -300.8 | 609.9 | 1.1 | 0.172 |
|  | -1.184 |  |  |  | 0.056 | + | + | + |  |  |  | 7 | -298.4 | 611.6 | 2.8 | 0.072 |
|  | -0.700 |  |  |  | 0.036 | + | + |  |  |  |  | 6 | -299.6 | 611.8 | 3.0 | 0.066 |
|  | -0.943 |  |  |  | 0.037 |  | + |  |  |  |  | 5 | -301.3 | 613.0 | 4.2 | 0.036 |
|  | -0.706 |  |  |  | 0.036 | + | + |  |  | + |  | 7 | -299.4 | 613.6 | 4.8 | 0.027 |
|  | -2.219 |  |  | 0.028 | 0.035 |  |  |  |  |  |  | 5 | -302.2 | 614.8 | 6.0 | 0.015 |
|  | -1.690 |  |  | 0.020 | 0.035 | + |  |  |  |  |  | 6 | -301.7 | 615.8 | 7.0 | 0.009 |
|  | -2.013 |  |  | 0.016 | 0.054 | + |  | + |  |  |  | 7 | -300.9 | 616.4 | 7.6 | 0.006 |
|  | -2.257 |  |  | 0.028 | 0.035 |  | + |  |  |  |  | 6 | -302.7 | 617.9 | 9.1 | 0.003 |
|  | -0.778 | 0.001 |  |  | 0.037 | + |  |  |  |  |  | 6 | -303.1 | 618.8 | 10.0 | 0.002 |
|  | -1.323 | 0.003 |  |  | 0.056 | + |  | + |  |  |  | 7 | -302.0 | 618.8 | 10.0 | 0.002 |
|  | -1.645 |  |  | 0.019 | 0.035 | + | + |  |  |  |  | 7 | -302.1 | 619.0 | 10.2 | 0.002 |
|  | -0.686 |  | -0.003 |  | 0.036 | + |  |  |  |  |  | 6 | -303.3 | 619.1 | 10.3 | 0.002 |
|  | -1.179 |  | -0.003 |  | 0.055 | + |  | + |  |  |  | 7 | -302.2 | 619.2 | 10.4 | 0.002 |
|  | -1.024 | 0.003 |  |  | 0.037 |  |  |  |  |  |  | 5 | -304.6 | 619.7 | 10.9 | 0.001 |
|  | -0.822 |  | -0.005 |  | 0.036 |  |  |  |  |  |  | 5 | -304.7 | 619.7 | 10.9 | 0.001 |
|  | -0.687 |  |  |  | 0.036 | + | + |  | + |  |  | 7 | -303.0 | 620.7 | 11.9 | 0.001 |
|  | -0.696 | 0.000 |  |  | 0.036 | + | + |  |  |  |  | 7 | -303.5 | 621.7 | 12.9 | 0.000 |
|  | -0.900 |  |  |  | 0.035 |  | + |  | + |  |  | 6 | -304.6 | 621.8 | 13.0 | 0.000 |
|  | -0.607 |  | -0.003 |  | 0.035 | + | + |  |  |  |  | 7 | -303.7 | 622.1 | 13.2 | 0.000 |
|  | -0.770 |  | -0.006 |  | 0.035 |  | + |  |  |  |  | 6 | -305.1 | 622.7 | 13.9 | 0.000 |
|  | -1.011 | 0.003 |  |  | 0.037 |  | + |  |  |  |  | 6 | -305.1 | 622.8 | 14.0 | 0.000 |
|  | -2.344 | -0.006 |  | 0.033 | 0.035 |  |  |  |  |  |  | 6 | -305.7 | 624.0 | 15.2 | 0.000 |
|  | -2.166 |  | -0.001 | 0.027 | 0.035 |  |  |  |  |  |  | 6 | -306.3 | 625.2 | 16.4 | 0.000 |
|  | -1.812 | -0.005 |  | 0.024 | 0.035 | + |  |  |  |  |  | 7 | -305.3 | 625.3 | 16.5 | 0.000 |
|  | -1.672 |  | 0.000 | 0.019 | 0.035 | + |  |  |  |  |  | 7 | -305.8 | 626.3 | 17.5 | 0.000 |
|  | 0.214 |  |  |  |  | + |  |  |  |  |  | 4 | -309.0 | 626.3 | 17.5 | 0.000 |
|  | -2.234 |  |  | 0.028 | 0.035 |  | + |  | + |  |  | 7 | -306.0 | 626.8 | 18.0 | 0.000 |
|  | -2.344 | -0.006 |  | 0.033 | 0.035 |  | + |  |  |  |  | 7 | -306.2 | 627.2 | 18.4 | 0.000 |
|  | -2.218 |  | 0.000 | 0.028 | 0.035 |  | + |  |  |  |  | 7 | -306.8 | 628.3 | 19.5 | 0.000 |
|  | 0.298 |  |  |  |  | + | + |  |  |  |  | 5 | -309.2 | 628.8 | 20.0 | 0.000 |
|  | -0.722 | 0.002 | -0.003 |  | 0.036 | + |  |  |  |  |  | 7 | -307.1 | 629.0 | 20.2 | 0.000 |
|  | -0.895 | 0.005 | -0.006 |  | 0.036 |  |  |  |  |  |  | 6 | -308.3 | 629.2 | 20.4 | 0.000 |
|  | 0.247 |  |  |  |  | + | + |  |  | + |  | 6 | -308.8 | 630.1 | 21.3 | 0.000 |
|  | -1.730 |  |  | 0.037 |  |  |  |  |  |  |  | 4 | -311.0 | 630.2 | 21.4 | 0.000 |
|  | 0.047 |  |  |  |  |  | + |  |  |  |  | 4 | -311.1 | 630.4 | 21.6 | 0.000 |
|  | -0.725 |  | -0.006 |  | 0.034 |  | + |  | + |  |  | 7 | -308.4 | 631.5 | 22.7 | 0.000 |
|  | -0.968 | 0.003 |  |  | 0.035 |  | + |  | + |  |  | 7 | -308.4 | 631.6 | 22.8 | 0.000 |
|  | -1.233 |  |  | 0.029 |  | + |  |  |  |  |  | 5 | -310.6 | 631.6 | 22.8 | 0.000 |
|  | -0.857 | 0.005 | -0.006 |  | 0.035 |  | + |  |  |  |  | 7 | -308.8 | 632.3 | 23.5 | 0.000 |
|  | -1.731 |  |  | 0.037 |  |  | + |  |  |  |  | 5 | -311.4 | 633.2 | 24.4 | 0.000 |
|  | -2.541 | -0.008 | 0.002 | 0.037 | 0.035 |  |  |  |  |  |  | 7 | -309.7 | 634.0 | 25.2 | 0.000 |
|  | -1.141 |  |  | 0.028 |  | + | + |  |  |  |  | 6 | -311.0 | 634.5 | 25.7 | 0.000 |
|  | 0.381 |  | -0.009 |  |  | + |  |  |  |  |  | 5 | -312.1 | 634.6 | 25.8 | 0.000 |
|  | 0.245 |  | -0.011 |  |  |  |  |  |  |  |  | 4 | -313.2 | 634.7 | 25.9 | 0.000 |
|  | -1.150 |  |  | 0.028 |  | + | + |  |  | + |  | 7 | -310.6 | 636.0 | 27.2 | 0.000 |
|  | 0.202 | 0.001 |  |  |  | + |  |  |  |  |  | 5 | -312.9 | 636.1 | 27.3 | 0.000 |
|  | 0.508 |  | -0.010 |  |  | + | + |  |  |  |  | 6 | -312.1 | 636.8 | 27.9 | 0.000 |
|  | 0.342 |  | -0.012 |  |  |  | + |  |  |  |  | 5 | -313.4 | 637.1 | 28.3 | 0.000 |
|  | -0.053 | 0.003 |  |  |  |  |  |  |  |  |  | 4 | -314.5 | 637.2 | 28.4 | 0.000 |
|  | 0.465 |  | -0.010 |  |  | + | + |  |  | + |  | 7 | -311.7 | 638.2 | 29.4 | 0.000 |
|  | 0.340 | -0.002 |  |  |  | + | + |  |  |  |  | 6 | -313.0 | 638.5 | 29.7 | 0.000 |
|  | -1.941 | -0.010 |  | 0.046 |  |  |  |  |  |  |  | 5 | -314.2 | 638.7 | 29.9 | 0.000 |
|  | -1.310 |  | -0.006 | 0.031 |  |  |  |  |  |  |  | 5 | -314.6 | 639.6 | 30.8 | 0.000 |
|  | 0.287 | -0.002 |  |  |  | + | + |  |  | + |  | 7 | -312.5 | 639.8 | 31.0 | 0.000 |
|  | 0.008 | 0.002 |  |  |  |  | + |  |  |  |  | 5 | -314.8 | 640.0 | 31.2 | 0.000 |
|  | -1.455 | -0.009 |  | 0.038 |  | + |  |  |  |  |  | 6 | -313.9 | 640.3 | 31.5 | 0.000 |
|  | -0.858 |  | -0.006 | 0.024 |  | + |  |  |  |  |  | 6 | -314.3 | 641.2 | 32.3 | 0.000 |
|  | -1.891 | -0.011 |  | 0.046 |  |  | + |  |  |  |  | 6 | -314.5 | 641.6 | 32.8 | 0.000 |
|  | -1.215 |  | -0.007 | 0.030 |  |  | + |  |  |  |  | 6 | -315.0 | 642.5 | 33.7 | 0.000 |
|  | -1.309 | -0.010 |  | 0.036 |  | + | + |  |  |  |  | 7 | -314.2 | 643.0 | 34.2 | 0.000 |
|  | 0.137 | 0.007 | -0.013 |  |  |  |  |  |  |  |  | 5 | -316.7 | 643.8 | 35.0 | 0.000 |
|  | -0.615 |  | -0.007 | 0.021 |  | + | + |  |  |  |  | 7 | -314.6 | 643.8 | 35.0 | 0.000 |
|  | 0.307 | 0.004 | -0.010 |  |  | + |  |  |  |  |  | 6 | -315.8 | 644.1 | 35.3 | 0.000 |
|  | 0.465 | 0.002 | -0.010 |  |  | + | + |  |  |  |  | 7 | -315.8 | 646.4 | 37.6 | 0.000 |
|  | 0.235 | 0.006 | -0.013 |  |  |  | + |  |  |  |  | 6 | -316.9 | 646.4 | 37.6 | 0.000 |
|  | -1.638 | -0.007 | -0.004 | 0.040 |  |  |  |  |  |  |  | 6 | -318.0 | 648.5 | 39.7 | 0.000 |
|  | -1.159 | -0.006 | -0.004 | 0.032 |  | + |  |  |  |  |  | 7 | -317.7 | 650.1 | 41.3 | 0.000 |
|  | -1.536 | -0.008 | -0.004 | 0.039 |  |  | + |  |  |  |  | 7 | -318.3 | 651.3 | 42.5 | 0.000 |
| *Exploratory Behavior* | -0.878 |  |  |  |  | + | + |  |  |  |  | 5 | -240.9 | 492.4 | 0.0 | 0.379 |
|  | -0.828 |  |  |  |  | + | + |  |  | + |  | 6 | -240.0 | 492.8 | 0.3 | 0.319 |
|  | -0.481 |  |  |  |  | + |  |  |  |  |  | 4 | -242.9 | 494.2 | 1.7 | 0.159 |
|  | -0.472 |  |  |  | -0.023 | + | + |  |  |  |  | 6 | -242.4 | 497.5 | 5.1 | 0.029 |
|  | -0.408 |  |  |  | -0.023 | + | + |  |  | + |  | 7 | -241.4 | 497.9 | 5.4 | 0.025 |
|  | -1.267 |  | 0.023 |  |  | + | + |  |  |  |  | 6 | -243.0 | 498.7 | 6.3 | 0.016 |
|  | -1.229 |  | 0.023 |  |  | + | + |  |  | + |  | 7 | -242.1 | 499.2 | 6.7 | 0.013 |
|  | -0.071 |  |  |  | -0.022 | + |  |  |  |  |  | 5 | -244.5 | 499.5 | 7.1 | 0.011 |
|  | -0.780 |  |  | -0.002 |  | + | + |  |  |  |  | 6 | -243.7 | 500.1 | 7.6 | 0.008 |
|  | -0.728 |  |  | -0.002 |  | + | + |  |  | + |  | 7 | -242.7 | 500.4 | 8.0 | 0.007 |
|  | -1.135 | 0.010 |  |  |  | + | + |  |  |  |  | 6 | -244.0 | 500.7 | 8.3 | 0.006 |
|  | -0.221 |  |  |  |  |  | + |  |  |  |  | 4 | -246.3 | 501.0 | 8.6 | 0.005 |
|  | -1.087 | 0.010 |  |  |  | + | + |  |  | + |  | 7 | -243.0 | 501.1 | 8.6 | 0.005 |
|  | -0.732 |  | 0.017 |  |  | + |  |  |  |  |  | 5 | -245.5 | 501.5 | 9.1 | 0.004 |
|  | 0.207 |  |  | -0.014 |  | + |  |  |  |  |  | 5 | -245.5 | 501.5 | 9.1 | 0.004 |
|  | -0.438 | -0.002 |  |  |  | + |  |  |  |  |  | 5 | -246.2 | 502.9 | 10.5 | 0.002 |
|  | -0.606 |  |  |  | -0.016 | + | + | + |  |  |  | 7 | -244.8 | 504.6 | 12.2 | 0.001 |
|  | -0.967 |  |  | 0.010 | -0.024 | + | + |  |  |  |  | 7 | -245.0 | 505.1 | 12.6 | 0.001 |
|  | -0.808 |  | 0.016 |  | -0.019 | + | + |  |  |  |  | 7 | -245.2 | 505.3 | 12.9 | 0.001 |
|  | -0.495 |  |  |  | -0.022 | + | + |  | + |  |  | 7 | -245.3 | 505.5 | 13.1 | 0.001 |
|  | 0.415 |  |  |  | -0.023 |  |  |  |  |  |  | 4 | -248.7 | 505.7 | 13.3 | 0.000 |
|  | -2.225 |  | 0.026 | 0.017 |  | + | + |  |  |  |  | 7 | -245.4 | 505.9 | 13.5 | 0.000 |
|  | 0.182 |  |  |  | -0.024 |  | + |  |  |  |  | 5 | -247.7 | 506.0 | 13.6 | 0.000 |
|  | -0.679 | 0.008 |  |  | -0.023 | + | + |  |  |  |  | 7 | -245.5 | 506.1 | 13.6 | 0.000 |
|  | 1.776 |  |  | -0.037 |  |  |  |  |  |  |  | 4 | -249.0 | 506.4 | 14.0 | 0.000 |
|  | -0.193 |  |  |  | -0.016 | + |  | + |  |  |  | 6 | -247.0 | 506.7 | 14.2 | 0.000 |
|  | -0.719 |  | 0.027 |  |  |  | + |  |  |  |  | 5 | -248.1 | 506.7 | 14.3 | 0.000 |
|  | -0.360 |  | 0.023 |  |  |  |  |  |  |  |  | 4 | -249.4 | 507.1 | 14.7 | 0.000 |
|  | 0.122 |  |  | -0.004 | -0.022 | + |  |  |  |  |  | 6 | -247.2 | 507.1 | 14.7 | 0.000 |
|  | 1.369 |  |  | -0.033 |  |  | + |  |  |  |  | 5 | -248.3 | 507.2 | 14.7 | 0.000 |
|  | -1.258 | -0.001 | 0.023 |  |  | + | + |  |  |  |  | 7 | -246.1 | 507.3 | 14.9 | 0.000 |
|  | -0.262 |  | 0.010 |  | -0.020 | + |  |  |  |  |  | 6 | -247.6 | 507.9 | 15.4 | 0.000 |
|  | -0.555 | 0.014 |  | -0.013 |  | + | + |  |  |  |  | 7 | -246.5 | 507.9 | 15.5 | 0.000 |
|  | 0.024 | -0.004 |  |  | -0.023 | + |  |  |  |  |  | 6 | -247.8 | 508.3 | 15.8 | 0.000 |
|  | 0.159 | -0.008 |  |  |  |  |  |  |  |  |  | 4 | -250.3 | 508.9 | 16.5 | 0.000 |
|  | -0.549 |  | 0.017 | -0.004 |  | + |  |  |  |  |  | 6 | -248.1 | 509.0 | 16.6 | 0.000 |
|  | -0.193 | -0.001 |  |  |  |  | + |  |  |  |  | 5 | -249.4 | 509.4 | 17.0 | 0.000 |
|  | -0.550 | -0.013 | 0.024 |  |  | + |  |  |  |  |  | 6 | -248.4 | 509.5 | 17.1 | 0.000 |
|  | 0.304 | 0.003 |  | -0.017 |  | + |  |  |  |  |  | 6 | -248.6 | 510.0 | 17.6 | 0.000 |
|  | 1.707 |  |  | -0.028 | -0.021 |  |  |  |  |  |  | 5 | -250.9 | 512.3 | 19.8 | 0.000 |
|  | 1.245 |  |  | -0.022 | -0.022 |  | + |  |  |  |  | 6 | -250.1 | 512.9 | 20.4 | 0.000 |
|  | -0.250 |  | 0.020 |  | -0.020 |  | + |  |  |  |  | 6 | -250.2 | 513.2 | 20.7 | 0.000 |
|  | 0.111 |  | 0.015 |  | -0.020 |  |  |  |  |  |  | 5 | -251.5 | 513.4 | 21.0 | 0.000 |
|  | 0.945 |  | 0.018 | -0.026 |  |  |  |  |  |  |  | 5 | -251.6 | 513.7 | 21.3 | 0.000 |
|  | 0.097 |  | 0.024 | -0.016 |  |  | + |  |  |  |  | 6 | -250.5 | 513.7 | 21.3 | 0.000 |
|  | 0.194 |  |  |  | -0.024 |  | + |  | + |  |  | 6 | -250.6 | 513.9 | 21.5 | 0.000 |
|  | -0.081 | -0.022 | 0.032 |  |  |  |  |  |  |  |  | 5 | -251.7 | 513.9 | 21.5 | 0.000 |
|  | 0.616 | -0.010 |  |  | -0.024 |  |  |  |  |  |  | 5 | -251.7 | 514.0 | 21.5 | 0.000 |
|  | -0.461 | -0.016 | 0.033 |  |  |  | + |  |  |  |  | 6 | -250.8 | 514.3 | 21.9 | 0.000 |
|  | 0.052 |  |  | -0.005 | -0.016 | + |  | + |  |  |  | 7 | -249.7 | 514.3 | 21.9 | 0.000 |
|  | 0.262 | -0.003 |  |  | -0.024 |  | + |  |  |  |  | 6 | -250.9 | 514.5 | 22.0 | 0.000 |
|  | 1.904 | 0.004 |  | -0.042 |  |  |  |  |  |  |  | 5 | -252.0 | 514.6 | 22.2 | 0.000 |
|  | 1.577 | 0.011 |  | -0.042 |  |  | + |  |  |  |  | 6 | -251.1 | 515.0 | 22.6 | 0.000 |
|  | -0.381 |  | 0.010 |  | -0.014 | + |  | + |  |  |  | 7 | -250.0 | 515.1 | 22.7 | 0.000 |
|  | -0.329 |  | 0.011 | 0.001 | -0.020 | + |  |  |  |  |  | 7 | -250.2 | 515.4 | 23.0 | 0.000 |
|  | -2.187 | -0.030 | 0.037 | 0.036 |  | + |  |  |  |  |  | 7 | -250.2 | 515.4 | 23.0 | 0.000 |
|  | -0.091 | -0.005 |  |  | -0.016 | + |  | + |  |  |  | 7 | -250.2 | 515.5 | 23.0 | 0.000 |
|  | 0.034 | -0.004 |  | 0.000 | -0.023 | + |  |  |  |  |  | 7 | -250.3 | 515.6 | 23.1 | 0.000 |
|  | -0.114 | -0.012 | 0.016 |  | -0.020 | + |  |  |  |  |  | 7 | -250.5 | 516.0 | 23.6 | 0.000 |
|  | -0.554 | -0.028 | 0.036 | 0.011 |  |  |  |  |  |  |  | 6 | -253.8 | 520.3 | 27.9 | 0.000 |
|  | 1.174 |  | 0.012 | -0.021 | -0.020 |  |  |  |  |  |  | 6 | -253.8 | 520.3 | 27.9 | 0.000 |
|  | 0.323 |  | 0.018 | -0.011 | -0.020 |  | + |  |  |  |  | 7 | -252.7 | 520.3 | 27.9 | 0.000 |
|  | 0.351 | -0.021 | 0.025 |  | -0.020 |  |  |  |  |  |  | 6 | -253.9 | 520.5 | 28.1 | 0.000 |
|  | 1.674 | -0.002 |  | -0.026 | -0.022 |  |  |  |  |  |  | 6 | -253.9 | 520.5 | 28.1 | 0.000 |
|  | -0.969 | -0.021 | 0.037 | 0.011 |  |  | + |  |  |  |  | 7 | -252.8 | 520.7 | 28.2 | 0.000 |
|  | 1.247 |  |  | -0.022 | -0.023 |  | + |  | + |  |  | 7 | -252.9 | 520.8 | 28.4 | 0.000 |
|  | -0.028 | -0.014 | 0.025 |  | -0.019 |  | + |  |  |  |  | 7 | -253.0 | 520.9 | 28.5 | 0.000 |
|  | 1.347 | 0.005 |  | -0.027 | -0.022 |  | + |  |  |  |  | 7 | -253.0 | 521.0 | 28.6 | 0.000 |
|  | -0.242 |  | 0.020 |  | -0.020 |  | + |  | + |  |  | 7 | -253.1 | 521.1 | 28.7 | 0.000 |
|  | 0.279 | -0.003 |  |  | -0.025 |  | + |  | + |  |  | 7 | -253.7 | 522.4 | 30.0 | 0.000 |
|  | -0.389 | -0.029 | 0.031 | 0.017 | -0.020 |  |  |  |  |  |  | 7 | -255.9 | 526.8 | 34.4 | 0.000 |
| *Feeding Accuracy* | 1.102 |  |  |  |  | + |  |  |  |  |  | 4 | 15.2 | -22.0 | 0.0 | 0.726 |
|  | 1.041 |  |  |  |  |  | + |  |  |  |  | 4 | 14.0 | -19.5 | 2.5 | 0.211 |
|  | 1.083 |  |  |  |  | + | + |  |  |  |  | 5 | 13.4 | -16.2 | 5.8 | 0.039 |
|  | 1.049 |  |  | 0.000 |  |  |  |  |  |  |  | 4 | 10.2 | -12.1 | 9.9 | 0.005 |
|  | 1.049 | 0.001 |  |  |  |  |  |  |  |  |  | 4 | 9.8 | -11.2 | 10.8 | 0.003 |
|  | 1.075 |  |  |  |  | + | + |  |  | + |  | 6 | 11.9 | -11.0 | 11.0 | 0.003 |
|  | 1.089 |  |  |  | -0.001 |  |  |  |  |  |  | 4 | 9.6 | -10.9 | 11.1 | 0.003 |
|  | 1.045 |  | 0.001 |  |  |  |  |  |  |  |  | 4 | 9.5 | -10.6 | 11.5 | 0.002 |
|  | 1.078 |  | 0.001 |  |  | + |  |  |  |  |  | 5 | 10.4 | -10.3 | 11.7 | 0.002 |
|  | 1.180 |  |  | -0.002 |  | + |  |  |  |  |  | 5 | 10.4 | -10.2 | 11.8 | 0.002 |
|  | 1.092 | 0.001 |  |  |  | + |  |  |  |  |  | 5 | 9.6 | -8.7 | 13.3 | 0.001 |
|  | 1.114 |  |  |  | 0.000 | + |  |  |  |  |  | 5 | 9.2 | -7.9 | 14.2 | 0.001 |
|  | 0.991 |  |  | 0.001 |  |  | + |  |  |  |  | 5 | 9.1 | -7.5 | 14.5 | 0.001 |
|  | 1.017 | 0.001 |  |  |  |  | + |  |  |  |  | 5 | 8.7 | -6.9 | 15.1 | 0.000 |
|  | 1.011 |  | 0.001 |  |  |  | + |  |  |  |  | 5 | 8.6 | -6.6 | 15.4 | 0.000 |
|  | 1.065 |  |  |  | -0.001 |  | + |  |  |  |  | 5 | 8.2 | -5.9 | 16.1 | 0.000 |
|  | 1.049 |  | 0.002 |  |  | + | + |  |  |  |  | 6 | 8.9 | -5.1 | 17.0 | 0.000 |
|  | 1.131 |  |  | -0.001 |  | + | + |  |  |  |  | 6 | 8.5 | -4.2 | 17.8 | 0.000 |
|  | 1.064 | 0.001 |  |  |  | + | + |  |  |  |  | 6 | 8.0 | -3.1 | 18.9 | 0.000 |
|  | 1.094 |  |  |  | 0.000 | + | + |  |  |  |  | 6 | 7.4 | -1.9 | 20.1 | 0.000 |
|  | 1.038 |  |  |  | 0.003 | + |  | + |  |  |  | 6 | 6.6 | -0.3 | 21.7 | 0.000 |
|  | 1.042 |  | 0.002 |  |  | + | + |  |  | + |  | 7 | 7.5 | 0.2 | 22.2 | 0.000 |
|  | 1.080 | 0.001 |  | -0.001 |  |  |  |  |  |  |  | 5 | 5.0 | 0.6 | 22.6 | 0.000 |
|  | 1.124 |  |  | -0.001 |  | + | + |  |  | + |  | 7 | 7.1 | 1.0 | 23.0 | 0.000 |
|  | 1.102 |  |  | 0.000 | -0.001 |  |  |  |  |  |  | 5 | 4.7 | 1.3 | 23.3 | 0.000 |
|  | 0.989 |  | 0.001 | 0.001 |  |  |  |  |  |  |  | 5 | 4.6 | 1.4 | 23.4 | 0.000 |
|  | 1.108 |  | 0.001 | -0.001 |  | + |  |  |  |  |  | 6 | 5.5 | 1.8 | 23.8 | 0.000 |
|  | 1.054 | 0.001 |  |  |  | + | + |  |  | + |  | 7 | 6.5 | 2.0 | 24.1 | 0.000 |
|  | 1.225 | 0.001 |  | -0.003 |  | + |  |  |  |  |  | 6 | 5.3 | 2.3 | 24.3 | 0.000 |
|  | 1.080 | 0.000 |  |  | -0.001 |  |  |  |  |  |  | 5 | 4.1 | 2.4 | 24.4 | 0.000 |
|  | 1.095 |  |  |  | -0.002 |  | + |  | + |  |  | 6 | 5.2 | 2.4 | 24.4 | 0.000 |
|  | 1.038 | 0.001 | 0.001 |  |  |  |  |  |  |  |  | 5 | 3.9 | 2.7 | 24.7 | 0.000 |
|  | 1.071 |  | 0.001 |  | -0.001 |  |  |  |  |  |  | 5 | 3.9 | 2.8 | 24.9 | 0.000 |
|  | 1.082 | 0.000 | 0.001 |  |  | + |  |  |  |  |  | 6 | 4.9 | 3.1 | 25.1 | 0.000 |
|  | 1.086 |  |  |  | 0.000 | + | + |  |  | + |  | 7 | 6.0 | 3.2 | 25.2 | 0.000 |
|  | 1.206 |  |  | -0.002 | -0.001 | + |  |  |  |  |  | 6 | 4.5 | 3.8 | 25.9 | 0.000 |
|  | 1.087 |  | 0.001 |  | 0.000 | + |  |  |  |  |  | 6 | 4.4 | 4.0 | 26.1 | 0.000 |
|  | 0.880 |  | 0.001 | 0.003 |  |  | + |  |  |  |  | 6 | 4.1 | 4.6 | 26.7 | 0.000 |
|  | 1.027 | 0.001 |  | 0.000 |  |  | + |  |  |  |  | 6 | 3.9 | 5.0 | 27.0 | 0.000 |
|  | 1.106 | 0.000 |  |  | 0.000 | + |  |  |  |  |  | 6 | 3.7 | 5.5 | 27.5 | 0.000 |
|  | 1.021 |  |  |  | 0.003 | + | + | + |  |  |  | 7 | 4.6 | 5.8 | 27.8 | 0.000 |
|  | 1.039 |  |  | 0.001 | -0.001 |  | + |  |  |  |  | 6 | 3.3 | 6.2 | 28.3 | 0.000 |
|  | 1.124 |  |  |  | -0.001 | + | + |  | + |  |  | 7 | 4.4 | 6.4 | 28.4 | 0.000 |
|  | 0.998 | 0.001 | 0.001 |  |  |  | + |  |  |  |  | 6 | 3.2 | 6.4 | 28.5 | 0.000 |
|  | 1.017 |  | 0.002 | 0.001 |  | + | + |  |  |  |  | 7 | 4.1 | 7.0 | 29.0 | 0.000 |
|  | 1.042 | 0.001 |  |  | -0.001 |  | + |  |  |  |  | 6 | 2.9 | 7.1 | 29.1 | 0.000 |
|  | 1.034 |  | 0.001 |  | -0.001 |  | + |  |  |  |  | 6 | 2.9 | 7.1 | 29.1 | 0.000 |
|  | 1.174 | 0.002 |  | -0.002 |  | + | + |  |  |  |  | 7 | 3.4 | 8.2 | 30.2 | 0.000 |
|  | 1.048 | 0.000 | 0.002 |  |  | + | + |  |  |  |  | 7 | 3.4 | 8.4 | 30.4 | 0.000 |
|  | 1.058 |  | 0.002 |  | 0.000 | + | + |  |  |  |  | 7 | 2.9 | 9.4 | 31.4 | 0.000 |
|  | 1.156 |  |  | -0.001 | -0.001 | + | + |  |  |  |  | 7 | 2.6 | 10.0 | 32.0 | 0.000 |
|  | 1.179 |  |  | -0.003 | 0.003 | + |  | + |  |  |  | 7 | 2.2 | 10.7 | 32.7 | 0.000 |
|  | 1.075 | 0.001 |  |  | 0.000 | + | + |  |  |  |  | 7 | 1.9 | 11.3 | 33.3 | 0.000 |
|  | 1.017 |  | 0.001 |  | 0.003 | + |  | + |  |  |  | 7 | 1.5 | 12.1 | 34.1 | 0.000 |
|  | 1.043 | 0.000 |  |  | 0.003 | + |  | + |  |  |  | 7 | 1.0 | 13.1 | 35.1 | 0.000 |
|  | 0.997 | 0.000 | 0.001 | 0.001 |  |  |  |  |  |  |  | 6 | -0.7 | 14.1 | 36.2 | 0.000 |
|  | 1.118 | 0.001 |  | -0.001 | -0.001 |  |  |  |  |  |  | 6 | -0.7 | 14.2 | 36.2 | 0.000 |
|  | 1.097 |  |  | 0.000 | -0.002 |  | + |  | + |  |  | 7 | 0.3 | 14.6 | 36.6 | 0.000 |
|  | 1.104 | 0.000 | 0.001 | 0.000 |  | + |  |  |  |  |  | 7 | 0.2 | 14.7 | 36.7 | 0.000 |
|  | 1.043 |  | 0.001 | 0.001 | -0.001 |  |  |  |  |  |  | 6 | -1.0 | 14.9 | 36.9 | 0.000 |
|  | 1.088 | 0.000 |  |  | -0.002 |  | + |  | + |  |  | 7 | -0.3 | 15.7 | 37.7 | 0.000 |
|  | 1.067 |  | 0.001 |  | -0.002 |  | + |  | + |  |  | 7 | -0.3 | 15.8 | 37.8 | 0.000 |
|  | 1.129 |  | 0.001 | -0.001 | 0.000 | + |  |  |  |  |  | 7 | -0.5 | 16.1 | 38.1 | 0.000 |
|  | 1.070 | 0.000 | 0.001 |  | -0.001 |  |  |  |  |  |  | 6 | -1.7 | 16.1 | 38.2 | 0.000 |
|  | 1.237 | 0.001 |  | -0.003 | 0.000 | + |  |  |  |  |  | 7 | -0.7 | 16.5 | 38.5 | 0.000 |
|  | 1.096 | 0.000 | 0.001 |  | 0.000 | + |  |  |  |  |  | 7 | -1.1 | 17.3 | 39.3 | 0.000 |
|  | 0.873 | 0.000 | 0.001 | 0.003 |  |  | + |  |  |  |  | 7 | -1.2 | 17.5 | 39.5 | 0.000 |
|  | 0.924 |  | 0.001 | 0.002 | -0.001 |  | + |  |  |  |  | 7 | -1.8 | 18.7 | 40.7 | 0.000 |
|  | 1.059 | 0.001 |  | 0.000 | -0.001 |  | + |  |  |  |  | 7 | -1.9 | 19.0 | 41.0 | 0.000 |
|  | 1.025 | 0.000 | 0.001 |  | -0.001 |  | + |  |  |  |  | 7 | -2.6 | 20.4 | 42.4 | 0.000 |
|  | 1.031 | 0.000 | 0.001 | 0.001 | -0.001 |  |  |  |  |  |  | 7 | -6.3 | 27.7 | 49.7 | 0.000 |
| *Multivariate Phenotypic Differentiation* | 0.124 |  | 0.034 |  |  | + | + |  |  |  |  | 5 | -36.6 | 85.8 | 0.0 | 0.554 |
|  | -0.290 |  | 0.036 |  |  | + |  |  |  |  |  | 4 | -39.5 | 88.7 | 2.8 | 0.135 |
|  | 0.162 |  | 0.034 |  |  | + | + |  |  | + |  | 6 | -36.4 | 88.9 | 3.0 | 0.121 |
|  | -0.455 |  | 0.035 | 0.011 |  | + | + |  |  |  |  | 6 | -36.4 | 88.9 | 3.1 | 0.120 |
|  | -1.725 |  | 0.039 | 0.028 |  | + |  |  |  |  |  | 5 | -38.8 | 90.3 | 4.4 | 0.060 |
|  | 0.877 |  |  |  |  | + | + |  |  |  |  | 4 | -43.1 | 95.9 | 10.1 | 0.004 |
|  | 0.446 |  |  |  |  | + |  |  |  |  |  | 3 | -45.6 | 98.2 | 12.4 | 0.001 |
|  | 1.779 |  |  | -0.018 |  | + | + |  |  |  |  | 5 | -42.9 | 98.5 | 12.7 | 0.001 |
|  | -3.119 |  | 0.036 | 0.049 |  |  |  |  |  |  |  | 4 | -44.5 | 98.7 | 12.8 | 0.001 |
|  | -0.686 |  | 0.030 |  |  |  |  |  |  |  |  | 3 | -45.9 | 98.7 | 12.9 | 0.001 |
|  | -0.337 |  | 0.028 |  |  |  | + |  |  |  |  | 4 | -44.5 | 98.8 | 13.0 | 0.001 |
|  | 0.888 |  |  |  |  | + | + |  |  | + |  | 5 | -43.1 | 98.9 | 13.1 | 0.001 |
|  | -2.302 |  | 0.033 | 0.038 |  |  | + |  |  |  |  | 5 | -43.7 | 100.1 | 14.3 | 0.000 |
|  | 0.341 |  |  | 0.002 |  | + |  |  |  |  |  | 4 | -45.6 | 100.9 | 15.1 | 0.000 |
|  | 1.779 |  |  | -0.018 |  | + | + |  |  | + |  | 6 | -42.9 | 101.8 | 16.0 | 0.000 |
|  | 0.355 |  |  |  |  |  | + |  |  |  |  | 3 | -47.5 | 102.1 | 16.2 | 0.000 |
|  | -1.073 |  |  | 0.023 |  |  |  |  |  |  |  | 3 | -48.8 | 104.5 | 18.7 | 0.000 |
|  | -0.104 |  |  | 0.009 |  |  | + |  |  |  |  | 4 | -47.5 | 104.7 | 18.9 | 0.000 |
